# GABAergic amacrine cells balance biased chromatic information in the mouse retina

**DOI:** 10.1101/2024.03.08.584060

**Authors:** Maria M. Korympidou, Sarah Strauss, Timm Schubert, Katrin Franke, Philipp Berens, Thomas Euler, Anna Vlasits

**Author notes:** Equal authorship (Maria M. Korympidou, Sarah Strauss).

## Abstract

The retina extracts chromatic information present in an animal’s environment. In the mouse, the feed-forward, excitatory pathway through the retina is dominated by a chromatic gradient, with green and UV signals primarily processed in the dorsal and ventral retina, respectively. However, at the output of the retina, chromatic tuning is more mixed, suggesting that amacrine cells alter spectral tuning. We genetically targeted the population of 40+ GABAergic amacrine cell types and used two-photon calcium imaging to systematically survey chromatic responses in their dendritic processes. We found that amacrine cells show diverse chromatic responses in different spatial regions of their receptive fields and across the dorso-ventral axis of the retina. Compared to their excitatory inputs from bipolar cells, amacrine cells are less chromatically tuned and less likely to be colour-opponent. We identified 25 functional amacrine cell types that, in addition to their chromatic properties, exhibit distinctive achromatic receptive field properties. A combination of pharmacological interventions and a biologically-inspired deep learning model revealed how lateral inhibition and recurrent excitatory inputs shape chromatic properties of amacrine cells. Our data suggest that amacrine cells balance the strongly biased spectral tuning of excitation in the mouse retina and thereby support increased diversity in chromatic information of the retinal output.

## Introduction

The retina is a powerful image processor that extracts a variety of visual features present in the environment and sends them in parallel to the brain (Wässle, 2004). Among those features, chromatic information is particularly important for the animals’ behaviour and survival in their environmental niches and forms the neural basis of colour vision (Baden and Osorio, 2019; Gerl and Morris, 2008). Colour vision starts with light detection by different photoreceptor types with distinct spectral tuning, followed by downstream processing by neural circuits in the retina and the brain. Local comparisons of chromatic signals creates colour-opponency at different retinal layers, which has been studied in detail in both vertebrate and invertebrate species (Szatko et al., 2020; Thoreson and Dacey, 2019). However, there are also visual tasks where chromatic information is not wanted; for instance, motion vision is expected to function independent of an object’s spectral properties (Rosa et al., 2016). The retinal mechanisms that counterbalance strongly biased spectral tuning are far from understood.

A useful dichromatic model to explore such colour-related mechanisms is the house mouse (Mus musculus). Mice possess a particular opsin gradient that leads to nonuniform spectral sensitivity along the dorsal-ventral axis of the retina (Applebury et al., 2000; Röhlich et al., 1994). The mouse expresses three opsins, M-opsin, S-opsin, and rhodopsin. While S-cones exclusively express S-opsin (Haverkamp et al., 2005), M-cones co-express S- and M-opsins with the proportion of S-opsin increasing towards the retina’s ventral edge. In addition, the ventral retina contains an area in which S-cones are concentrated (Nadal-Nicolás et al., 2020). The result of this asymmetrical cone opsin distribution is a mostly green-sensitive dorsal and a mostly UV-sensitive ventral retina (Baden et al., 2013). Such a distribution should be detrimental to colour vision, because spectral comparisons would be difficult. Yet, colour-opponent signals have been reported in the mouse retina (Ekesten and Gouras, 2005; Joesch and Meister, 2016; Stabio et al., 2018; Szatko et al., 2020) and brain (Feord et al., 2023; Franke et al., 2023; Mouland et al., 2021; Rhim and Nauhaus, 2023), and mice are able to perform colour discrimination tasks (Denman et al., 2018; Jacobs et al., 2004).

Starting at the level of the photoreceptors and following the vertical excitatory pathway, colour-opponency appears to be a robust feature across populations of different retinal classes (Szatko et al., 2020). Indeed, several circuits supporting colour-opponent signals up to the level of RGCs have been described over the years, including cone-type selective circuits (Behrens et al., 2016; Breuninger et al., 2011; Haverkamp et al., 2005; Nadal-Nicolás et al., 2020; Stabio et al., 2018), cone-type unselective circuits (Chang et al., 2013) or rod-cone opponency (Joesch and Meister, 2016; Khani and Gollisch, 2021; Szatko et al., 2020). Interestingly, spectral tuning in RGCs appears to be more diverse than the photoreceptor and bipolar cell (BC) tuning, with RGC colour-opponency restricted to relatively few specific RGC types (Szatko et al., 2020). This suggests that chromatic signals are processed in the inner retina to allow for more diverse RGC tuning, pointing to the inhibitory network of amacrine cells (ACs) as a critical stage of “re-tuning” and “de-biasing” chromatic information.

ACs represent the largest and most diverse class of retinal neurons (Baden et al., 2018; Masland, 2012), counting more than 60 types (Li et al., 2024; Matsumoto et al., 2024; Yan et al., 2020). They shape spatial and temporal features of receptive fields in the inner retina, providing feedback, feedforward, and lateral inhibition to BCs, RGCs and other ACs, respectively (Diamond, 2017). This inhibition underlies important computations in the retina, for example, detection of motion direction (Euler et al., 2002; Vlasits et al., 2016) or segregating objects from background (Lin and Masland, 2006; Ölveczky et al., 2003). ACs use diverse neurotransmitters, primarily GABA or glycine, but also others. Most ACs lack axons and possess synaptic output sites on their dendrites. Thus, their synaptic inputs and outputs are not necessarily spatially segregated (Euler and Denk, 2001). This, together with their great morphological diversity (Helmstaedter et al., 2013), makes their study challenging. So far, only a few AC types have been functionally characterised in-depth.

Select ACs have been proposed to participate in chromatic processing in a variety of species (Chen and Li, 2012; Mills et al., 2014; Sher and DeVries, 2012). In the mouse, GABAergic ACs may mediate centre or surround components of colour-opponent RGC types (Chang et al., 2013; Joesch and Meister, 2016; Stabio et al., 2018; Szatko et al., 2020). In addition, ACs have been implicated in nonlinear chromatic integration (Khani and Gollisch, 2021). The role of ACs in chromatic processing is often inferred by pharmacological manipulations during experiments measuring activity of RGCs, because the relevant ACs have not yet been identified. Recently, a population-level study of ACs in the zebrafish retina revealed that most ACs feature spectrally-simple tuning, but that they contribute to the preservation of chromatic information in BCs that otherwise would be lost (Wang et al., 2023). This supports the idea that it is the ACs that can re-tune chromatic information in the inner retina.

In the present study, we systematically characterised chromatic responses of the GABAergic AC population and investigated their role in re-tuning chromatic information. Compared to BCs, which have a strong spectral drive, ACs are less strongly chromatically tuned and do not form a clear chromatic pattern along the retina’s dorso-ventral axis. We report 25 functional AC types with diverse chromatic identities, response polarities, RF organisations, and IPL stratifications, few of which are restricted to specific retinal locations. We used pharmacological interventions and a biology-inspired circuit model to study how chromatic signals in ACs arise.

## Results

### Recording the population responses of GABAergic amacrine cell dendrites to chromatic stimuli

To investigate how inhibition in the inner retina shapes chromatic signals, we performed two-photon imaging of AC dendrites in the *ex vivo* mouse retina. We used the transgenic mouse Gad2-IRES-Cre x Ai95D, which expresses the Ca^2+^ indicator GCaMP6f in GABAergic ACs and a small fraction of RGCs (Martersteck et al., 2017; Sonoda et al., 2020). This allowed us to simultaneously record responses from subcellular regions on AC processes throughout the entire inner plexiform layer (IPL; **Fig. 1a,b**). To characterise the colour preferences of ACs, we used a random noise stimulus designed to capture chromatic preferences across the potentially-large spatial scale of AC receptive fields (RFs): three concentric regions (centre, local surround, and far surround; 100, 300, 800 μm in diameter, respectively) that flickered independently (**Fig. 1c**). This stimulus was presented using either a UV or green LED in separate epochs (similar to Szatko et al., 2020). We defined regions of interest (ROIs) using a local correlation approach (Zhao et al., 2020), and restricted our analysis to ROIs located in the IPL, defined by the borders of the inner nuclear layer (INL) and the ganglion cell layer (GCL; **Fig. 1b,d,i**).

**Fig. 1.**
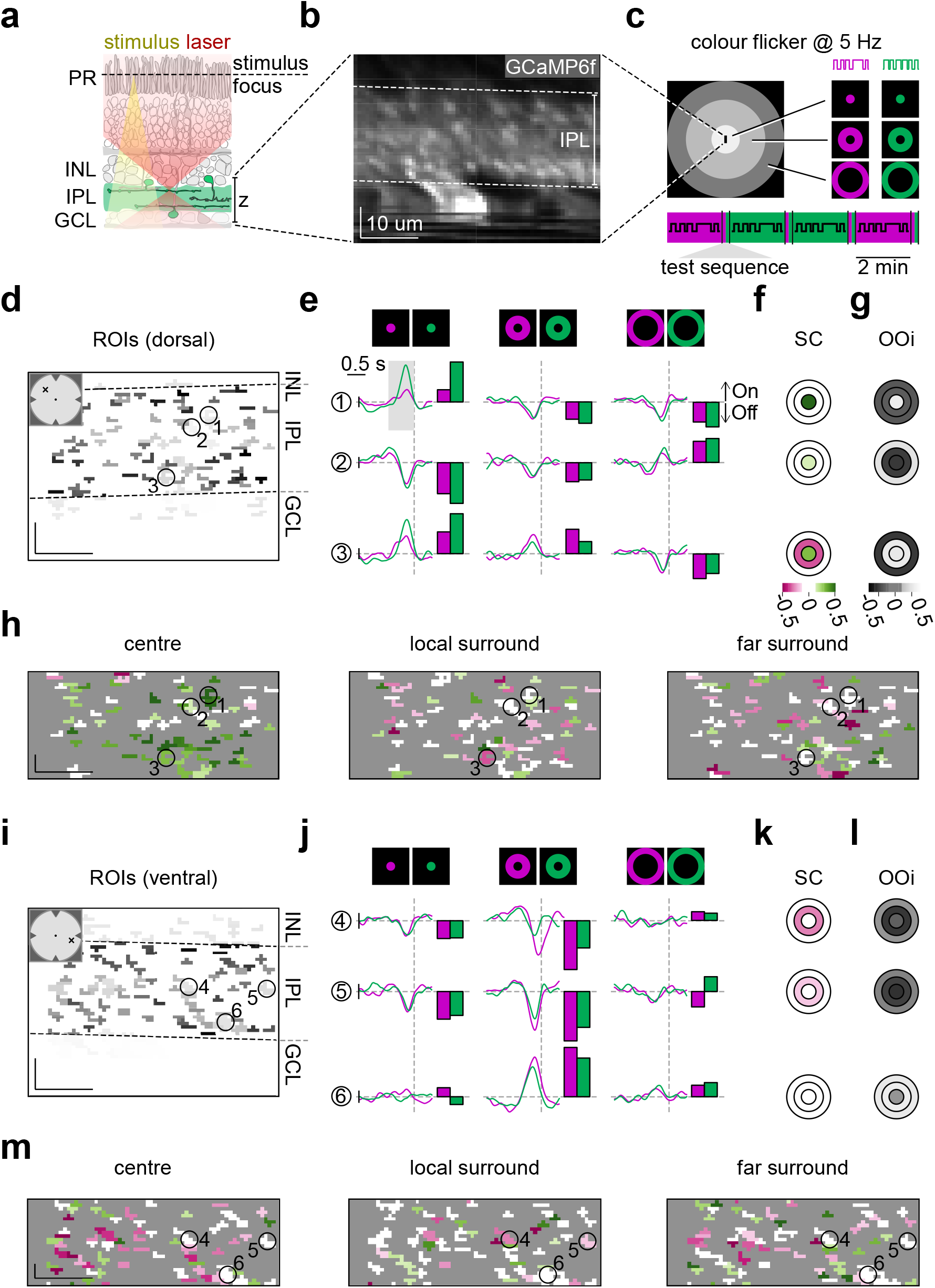
Diverse chromatic responses in amacrine cell processes. **(a)** Experimental setup used for amacrine cell (AC) recordings. PR, photoreceptors; INL, inner nuclear layer; IPL, inner plexiform layer; GCL, ganglion cell layer; red, yellow and green shadings represent two-photon excitation laser, stimulus, and GCaMP6f expression in GABAergic amacrine cells (Gad2-IRES-Cre x Ai95D mouse line), respectively. **(b)** Example x-z scan field (43 x 61 μm, 11.16 Hz) with GCaMP6f expression in the IPL. Dotted lines indicate borders with INL and GCL. Scale bars: 10 μm. **(c)** The chromatic stimulus consisted of 3 concentric regions – centre (100 μm in diameter, light grey), local surround (300 μm, dim grey), and far surround (800 μm, dark grey) – that were centred on the scan field and flickered randomly and independently for UV or green (see Methods). **(d)** Regions of interest (ROIs), indicating responsive AC processes, for an example scan field in the dorsal retina. **(f)** Chromatic preference of ROIs in centre, local surround, and far surround (cf. (c)) as estimated spectral contrast (*SC*). Green colour indicates a stronger preference for green stimulation, purple indicates a stronger preference for UV stimulation. **(g)** Like (f) but for response polarity as estimated On-Off index (*OOi*). Black indicates Off polarity, white indicates On polarity. **(h)** ROIs from (d) colour-coded according to ROIs’ *SC* for the three spatial conditions. **(i)** ROIs for an example scan field in the ventral retina (same field as in (b)). **(j-m)** Like in (e-h), but for ROIs shown in (i). In total, we recorded *n* = 79 scan fields in 17 mice.

To test the variation of AC chromatic signals across visual space, we collected Ca^2+^ responses from recording fields located in dorsal (**Fig. 1d-h**) and ventral retina (**Fig. 1i-m**). For each of the six stimulus conditions – centre, local surround, far surround, for both UV and green channels – we estimated a temporal kernel of the ROIs’ preferred stimulus (**Fig. 1e,j**). Subsequently, we estimated each ROI’s chromatic RF by calculating the spectral contrast index for each spatial stimulus condition (*SC*; *SC >* 0: green-preferring, *SC <* 0: UV-preferring; **Fig. 1f,h,k,m**). We found that AC RFs varied with respect to their *SC* across visual space, but also within each field (**Fig. 1h,m**). For example, ROI 1 preferred a green-biased centre stimulus but had similar preference for UV and green in the local and far surround, while ROI 3 also featured a green centre response but had a stronger preference for UV in the local surround (**Fig. 1e,f**).

In addition, we extracted the light polarity of each ROI, estimated as the On-Off index (*OOi*; *OOi >* 0: preferring increments of light, *OOi <* 0: preferring decrements of light; **Fig. 1g,l**). We also evaluated the polarity and temporal properties of each ROI using a “chirp” stimulus (see Methods and below). As expected, ROIs located closer to the GCL typically preferred increments of light (“On”, e.g., ROIs 3,6; but see ROI 1), while ROIs located closer to the INL usually preferred decrements of light (“Off”, e.g., ROIs 2,4,5). Thus, the sign of the *OOi* in the centre or local surround condition often matched the general On-Off subdivision of the IPL reported in the mammalian retina. These results show that our method allows us to systematically characterise the chromatic tuning and other response properties of AC processes across the retina and across the IPL.

### Chromatic tuning in amacrine cells is more diverse than in bipolar cells

Next, we explored whether the chromatic preferences of ACs follow the dorsal-ventral opsin gradient across the mouse retina, like the chromatic preferences of BCs, or whether the ACs have more mixed tuning like RGCs (Szatko et al., 2020). We recorded light-evoked Ca^2+^ responses from *n* = 4, 926 AC ROIs (*n* = 79 scan fields, *n* = 17 mice) distributed across the retina (**Fig. 2a**). To quantify how the AC chromatic tuning varied across retinal space, we divided the dorso-ventral axis into 500 μm bins. We found that many of the AC centre responses preferred UV and green stimulation equally, which we termed “achromatic”, whereas the rest were either green- or UV-tuned. We observed only slight preferences for green or UV with centre and local surround stimulation (**Fig. 2a,b**; “center”, “local surround”), while in the far surround preferences were quite mixed throughout the retina (**Fig. 2a,b**; “far surround”). Next, we examined spectral tuning as a function of location in the IPL. Notably, unlike in BCs, where there was stronger green tuning of the far surround in the Off layer in the ventral retina (Szatko et al., 2020), we did not observe any specific trend in AC chromatic preferences across different IPL layers (**Fig. 2c**).

**Fig. 2.**
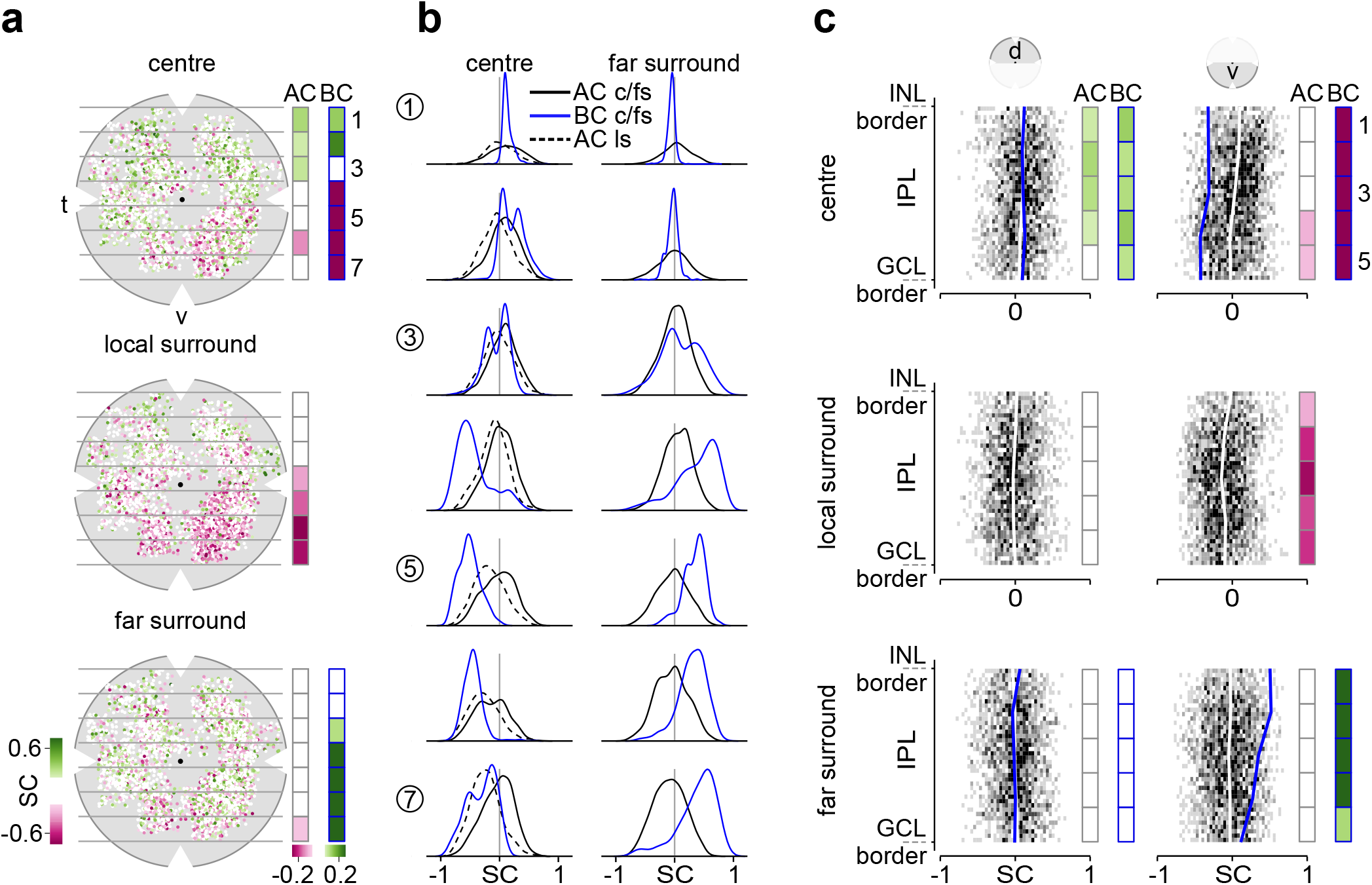
Chromatic tuning of amacrine cells differs from that of bipolar cells across the retina. **(a)** Amacrine cell (AC) chromatic preferences across the retina for responses to stimulus centre (top), local surround (middle), and far surround (bottom). Data points represent individual regions of interest (ROIs) and are colour-coded by spectral contrast (*SC*) (*n* = 4, 926 ROIs). For better visualisation, ROIs were scattered in x and y from the scan fields’ centre by *±* 412 μm. Gray horizontal lines and numbers on the right indicate arbitrarily defined bins across the retina (bin width: 0.57 mm). Right: Bars showing average *SC* per bin for ACs and bipolar cells (BCs) (for estimation of *SC* for *n* = 3, 270 BC ROIs from Szatko et al. (2020), see Methods). t, temporal; v, ventral. **(b)** *SC* distribution for ACs (black) and BCs (blue) from dorsal (bin 1) to ventral retina (bin 7) for centre (c), local surround (ls, dotted curves), and far surround (fs). Y axis represents density. **(c)** *SC* distribution across the inner plexiform layer (IPL) for dorsal (left, *n* = 2, 201 ROIs) and ventral (right, *n* = 2, 725 ROIs) AC ROIs and for centre (top), local surround (middle) and far surround (bottom). White and blue lines indicate mean *SC* across IPL for ACs and BCs (*n* = 1, 623 dorsal and *n* = 1, 647 ventral BC ROIs), respectively. Average *SC* for five equal bins across IPL depth is shown as bars for ACs and BCs (grey and blue outlines, respectively). INL, inner plexiform layer; GCL, ganglion cell layer. The local surround was not measured for BCs.

Next, we compared the AC chromatic preferences to the preferences of presynaptic BCs, which provide the excitatory drive to ACs, using a published dataset (Szatko et al., 2020). As described before, the tuning of BC centre responses follows the opsin expression gradient, resulting in green- and UV-dominant responses in dorsal and ventral retina, respectively (**Fig. 2a**; “center” BC bar). The tuning of BC surround responses, on the other hand, was shifted towards green in the ventral retina (**Fig. 2a**; “far surround” BC bar), resulting in an overall centre-surround colour-opponency in ventral BCs. In general, BC chromatic preferences were consistent at the population level (**Fig. 2b**). This was not the case in ACs, where AC chromatic preferences were much more broadly distributed and we did not find a marked centre-surround colour-opponency in ventral ACs (**Fig. 2a**; “center”, “far surround” AC bar)

In summary, we found that AC responses to chromatic stimuli are quite diverse and less spectrally tuned compared to BCs. Furthermore, AC chromatic tuning does not vary as strongly as BC tuning along the vertical axis of the retinal field. These results indicate that ACs may play a role in debiasing the strongly spectrally-tuned drive of BCs to RGCs and thereby, support the diverse chromatic tuning of RGCs.

### Identification of amacrine cell chromatic response types

Next, we sought to investigate AC chromatic identities across IPL layers and retinal regions. To cluster ROIs into functional types, we extracted features from each of the six AC temporal kernels separately using Principle Component Analysis (PCA) and then applied a Gaussian Mixture Model (GMM) (**Fig. 3a**; see also Baden et al., 2016). We identified 25 clusters, sorted them by median IPL depth and calculated their mean temporal kernels, *SC*, and *OOi* (**Fig. 3b-d**). Even though IPL depth was not used as a feature in our clustering, we found that most clusters were restricted to specific laminae within the IPL, which is in line with morphological data showing that most GABAergic ACs stratify narrowly in the IPL (**Fig. 3e**; see also Helmstaedter et al., 2013). However, there were also clusters that appeared to have representative ROIs in both On and Off IPL layers (e.g., *C*10, 11, 22, 24), which could correspond to bistratified ACs (Helmstaedter et al., 2013; Lin and Masland, 2006; Pérez De Sevilla Müller et al., 2007). We also found that individual clusters had stereotyped responses to the achromatic chirp stimulus, with clusters varying in their local vs. global response properties and their temporal kinetics (**Suppl. Fig. 2**).

**Fig. 3.**
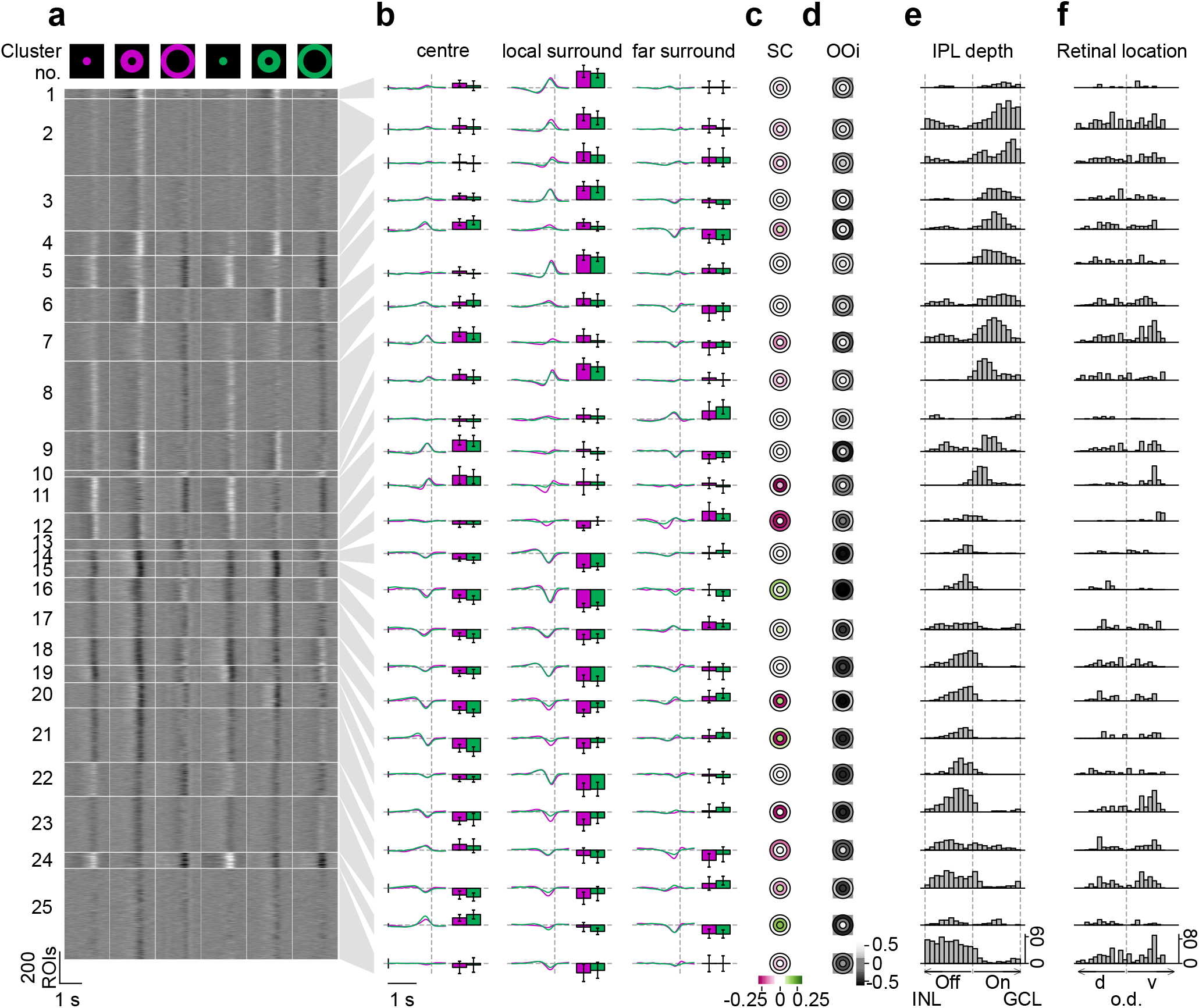
Functional characterisation of amacrine cells’ chromatic responses. **(a)** Temporal kernels calculated from the response of *n* = 5, 378 ROIs to UV and green, centre, local-surround, and far-surround flicker stimuli (cf. Fig. 1c), sorted into clusters. Clusters ordered according to their average inner plexiform layer (IPL) depth. Scale bars: x, 1 s; y, 200 ROIs. **(b)** Mean UV/green temporal kernels and colour tuning bars (*±*s.d.) of clusters for centre, local-surround, and far-surround. Vertical and horizontal dotted lines indicate response time and baseline, respectively. Scale bars: x, 1 s; y, 0.1 a.u. **(c)** Chromatic preference (as spectral contrast, *SC*, of mean cluster kernels) for centre (inner circle), local-surround (middle ring), and far-surround (outer ring) stimuli. **(d)** Like (c) but for response polarity (as On-Off index, *OOi*). **(e)** Distribution of cluster ROIs across IPL. Vertical dotted lines indicate borders to inner nuclear layer (INL), between On and Off IPL strata, and to ganglion cell layer (GCL), respectively. **(f)** Distribution of cluster ROIs along the retina’s dorso (d)–ventral (v) axis. ROIs for which the dorso-ventral location was not recorded are excluded from this plot (*n* = 452 ROIs). Dotted line indicates optic disc (o.d.) position.

Next, we examined the dorso-ventral locations of ROIs within each cluster (**Fig. 3f**). Here, we considered clusters with more than 80% of their ROIs in one retinal region to be region-specific and found three clusters satisfying this criterion (dorsal, *C*15; ventral, *C*12, *C*13). The remaining 22 clusters were “broad”, that is, their ROIs were more equally distributed between dorsal and ventral retina.

This lack of dorso-ventral organisation across AC clusters was in stark contrast with that reported for chromatic BC responses. To facilitate comparing AC and BC chromatic tuning, we performed the same clustering on the published BC dataset (Szatko et al., 2020). We identified 24 chromatic BC types (**Suppl. Fig. 1a,b**), which was roughly twice the number of genetically-identified BC types in the retina (Shekhar et al., 2016). The likely explanation for this discrepancy is that by including chromatic tuning into the clustering, BC types were “split” into two chromatic subtypes – depending on their position along the dorso-ventral axis. Indeed, BC clusters were confined to either the dorsal or the ventral retina (*n* = 10 ‘dorsal’ clusters; *n* = 13 ‘ventral’ clusters) (**Suppl. Fig. 1c**) and only one BC cluster (*BC*24) was broadly distributed. In addition, all BC clusters were chromatically tuned. In the dorsal retina, BCs were green-sensitive and non-colour opponent, whereas in the ventral retina, BCs were UV-sensitive and UV-green centre-surround opponent (**Suppl. Fig. 1d**). In addition, BC clusters were tightly stratified in IPL sublaminae and both dorsal and ventral retina comprised of On and Off clusters with similar chromatic tuning (**Suppl. Fig. 1e,f**).

Taken together, our data reveal a wide variety of functional chromatic AC types in the mouse retina and that there are distinct chromatic organisation principles for ACs and BCs.

### Achromatic features drive amacrine cell clustering into functional types

To better understand the organising principle of chromatic tuning in ACs, we performed hierarchical clustering of the cluster means (**Fig. 4a**). To our surprise, we found that clusters were first divided not by chromatic features, but by their achromatic centre kernel response polarity (*OOi*), and then by their spatial centre-surround RF organisation. We examined the spatial extent and the contrast preferences of AC RFs and identified families of clusters with common RF features.

**Fig. 4.**
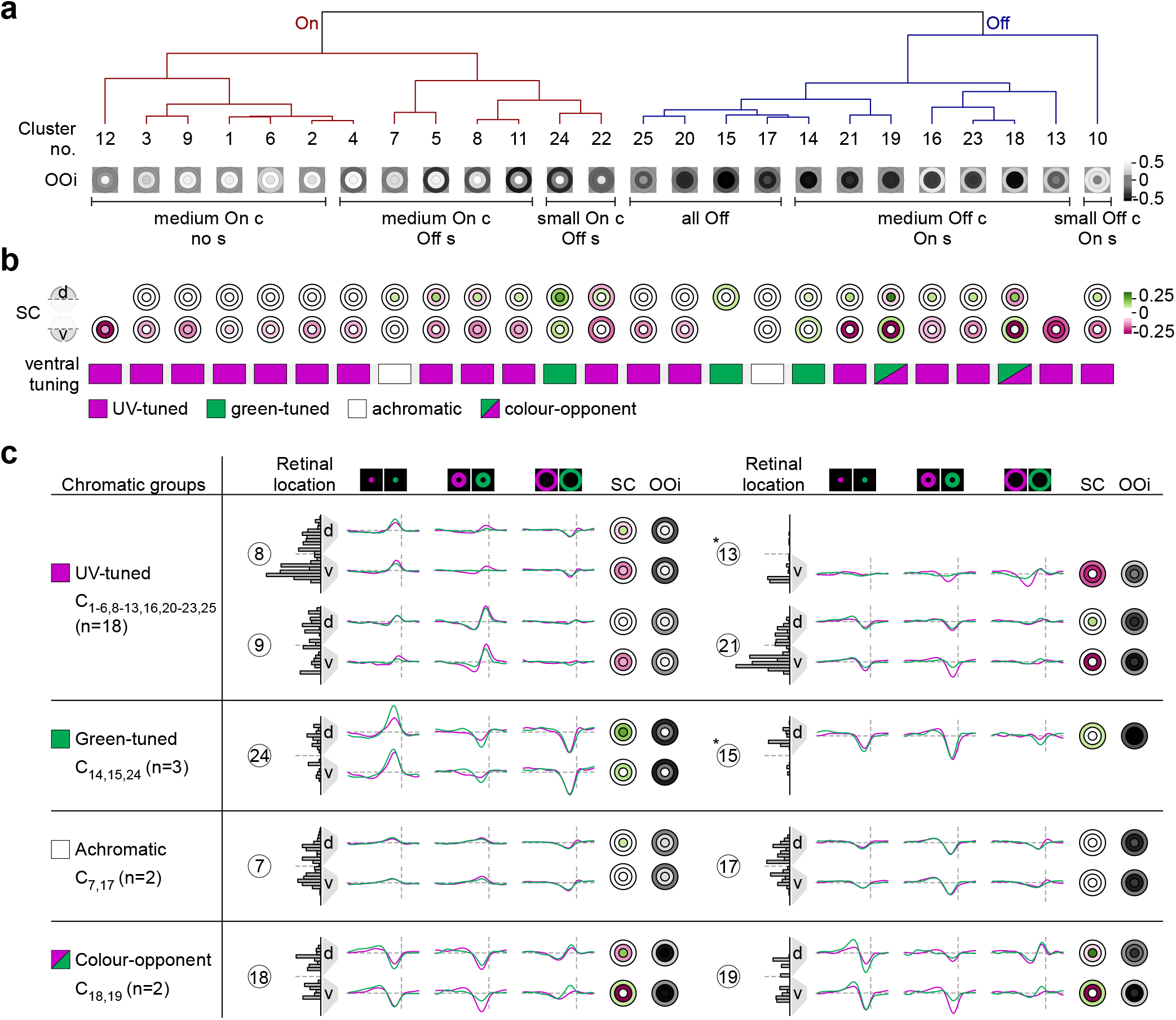
Amacrine cell functional clusters are largely driven by achromatic response properties. **(a)** Hierarchical clustering of amacrine cell (AC) clusters (cf. Fig. 3) based on kernel correlations. Clusters were divided into On (red) and Off (blue) groups, and further sorted according to their centre (c) vs. surround (s) receptive field (RF) organisation (for details, see text). *OOi* representations from Fig. 3d. **(b)** Chromatic preference of clusters’ representatives in dorsal (top) and ventral retina (bottom). Chromatic groups are named after the clusters’ average chromatic tuning in the ventral retina. **(c)** Example clusters from each chromatic group with member distribution, average kernels, chromatic preference and response polarity in dorsal and ventral retina. Asterisks indicate clusters with more than 80% of their members located in either dorsal or ventral retina. Retinal location plots from Fig. 3f.

One key difference was whether centre and local surround kernels were of opposite or same polarity, indicative of smaller (*≤*100 μm) or larger (between 100 and 300 μm) effective RF centre sizes, respectively. Both AC RF centre sizes were present in the On and Off branches of the hierarchy. Within these branches, the surround features drove the further subdivision into specific polarity groups. We observed that the surround (s) could be either non-responsive (“no s”), antagonistic (“On c/Off s” or “Off c/On s”) or same polarity (e.g., “all Off”) in relation to the centre and local surround (**Fig. 4a**). Within polarity groups, different chromatic types were quite, confirming that achromatic rather than chromatic response features dominated the AC clustering.

Beyond between-cluster chromatic differences, we also observed that the chromatic tuning within an individual AC cluster could be quite variable. Thus, we split each cluster into dorsal and ventral sub-populations and calculated the average *SC* separately (**Fig. 4b**). This analysis revealed that dorsal and ventral ROIs of a cluster can have different chromatic tuning on top of their shared contrast preferences (**Fig. 4b**). To categorise the chromatic tuning of AC clusters, we focused on the ventral retina, where more clusters were spectrally tuned. We identified four spectral groups based on the average cluster tuning: “UV-tuned” (*C*1 *−* 6, *C*8 *−* 13, *C*16, *C*20 *−* 23, *C*25), “green-tuned” (*C*14, *C*15, *C*24), “achromatic” (*C*7, *C*17), and “colour-opponent” (*C*18, *C*19) (**Fig. 4c**). ROIs in clusters *C*18 and *C*19 were colour-opponent in the ventral retina, showing centre-surround UV-Off/green-On responses and stratifying in the IPL’s Off sub-lamina. Several additional interesting RF motifs emerged from this analysis, which we review in the Discussion.

Taken together, we found that while ACs primarily cluster by their achromatic RF organisation and polarity, ACs also display diverse chromatic tuning, in particularly in the ventral retina. This indicates that ACs may support a range of chromatic operations in the inner retina.

### Inhibitory and excitatory contributions to GABAergic amacrine cells’ receptive fields

Given the complexity of AC chromatic tuning profiles compared to those of BCs, we next investigated how their tuning arises in the retinal circuit. First, we explored the role of inhibitory inputs from other ACs, which could be important in de-biasing chromatic input from BCs through feedforward inhibition. To test the role of interactions with other wide-field or narrow-field ACs (Diamond, 2017), we blocked ionotropic GABA (w/ gabazine+TPMPA) or glycine (w/ strychnine) receptors, respectively, while recording from GABAergic ACs (**Fig. 5a,e**). Blocking of GABA_A_ and GABA_C_ receptors decreased response amplitudes in both On and Off ACs across the retina. Surround responses switched polarity and most antagonistic RFs became non-antagonistic (**Fig. 5b left,c**). Notably, chromatic tuning of On and Off ROIs was affected mainly in the ventral retina, where the local- and far-surround chromatic preferences shifted towards UV (**Fig. 5b right,d**; see also **Suppl. Fig. 4a**). In contrast, disrupting glycinergic inhibition affected the response polarity less than blocking GABAergic inhibition and AC chromatic tuning was not altered significantly (**Fig. 5f-h**; see also **Suppl. Fig. 4b**). This suggests that lateral inhibition from GABAergic and/or glycinergic ACs mainly modulates GABAergic AC response polarity, while their chromatic properties rely to some extent on GABAergic inhibition. Under GABA receptor block, AC responses were more UV-shifted and, hence, became more similar to BC responses, especially in the ventral retina. This suggests that that GABAergic AC-AC interactions have a de-biasing function: they can counteract S-cone mediated UV dominance in the ventral retina.

**Fig. 5.**
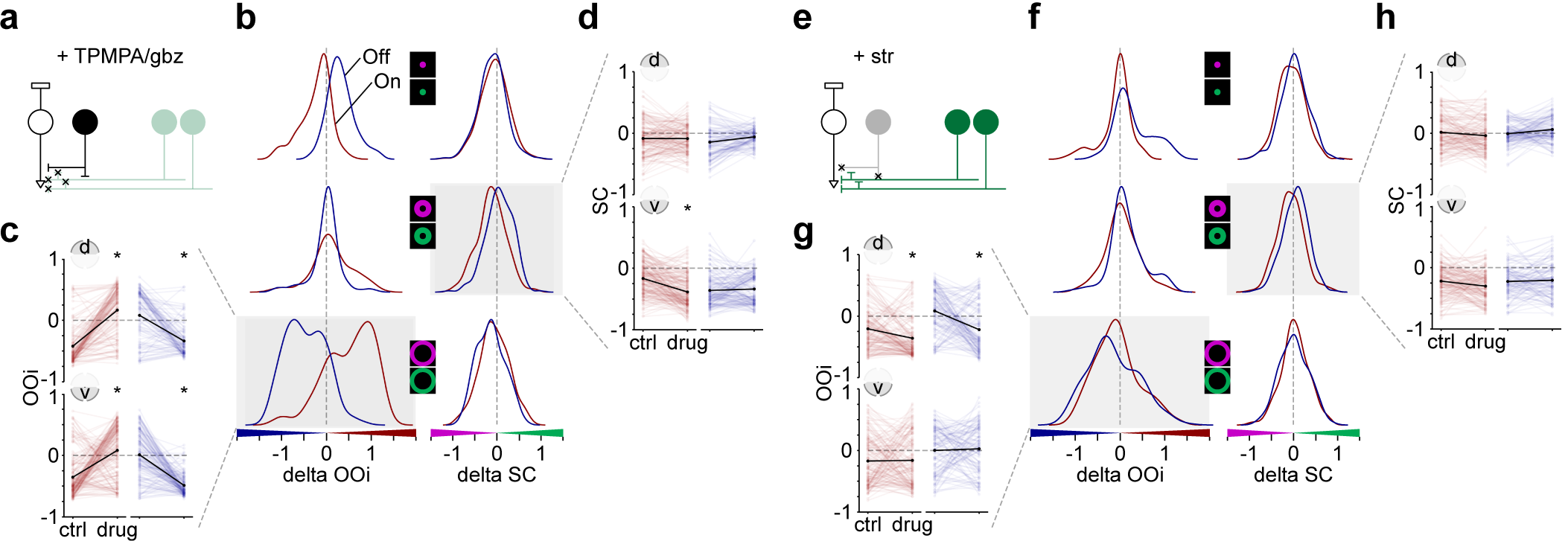
Effects of inhibition on amacrine cell chromatic responses. **(a)** In the inner retina, co-application of TPMPA (75 μM) and gabazine (gbz, 10 μM) removes ionotropic GABA receptor-mediated inhibition onto bipolar cells (BCs), small-field amacrine cells (ACs), and between wide-field ACs. Synapses onto retinal ganglion cells are not illustrated. TPMPA, (1,2,5,6-Tetrahydropyridin-4-yl)methylphosphinic acid. White, black, and green cells represent a BC, a small-field (glycinergic) AC, and wide-field (GABAergic) ACs, respectively. **(b)** Density plots of the difference in On-Off index (*delta OOi*) and spectral contrast (*delta SC*) between control and drug condition for centre (top), local-surround (middle) and far-surround (bottom) (*n* = 546 ROIs, *n* = 10 scan fields, *n* = 6 mice). We used the *OOi* of the “centre-control” condition to define On ROIs (*delta OOi >* 0, *n* = 310 ROIs, red) and Off ROIs (*delta OOi <* 0, *n* = 236 ROIs, blue). **(c)** Effect of TPMPA/gbz on far-surround *OOi*. ROIs separated into On (red) and Off (blue), as well as dorsal (top) and ventral (bottom). \**p <* 0.05, “ns” not significant; paired t-test corrected for multiple comparisons. **(d)** Effect of TPMPA/gbz on local-surround *SC*. Statistics same as in (c). **(e)** Strychnine (str, 0.5 μM) blocks glycine receptor-mediated inhibition onto Off BCs, wide-field ACs, and between small-field ACs. **(f-h)** Same as in (b-d) but for glycine receptor block (*n* = 497 (280 On, 217 Off) ROIs, *n* = 10 scan fields, *n* = 6 mice).

Next, we investigated whether the On and Off excitatory pathways had differential effects on AC chromatic responses. For this, we disrupted either the On or Off pathway by agonising metabotropic glutamate receptors (w/ L-AP4) or blocking kainate-sensitive ionotropic glutamate receptors (w/ UBP 310) to disrupt glutamate release from On and Off BCs, respectively (**Suppl. Fig. 3a,e**; see also Borghuis et al., 2014; Nakajima et al., 1993; Puller et al., 2013). When blocking the Off pathway, we observed that RF polarity was affected only in Off ACs of both dorsal and ventral retina, whereas chromatic tuning changed only for ventrally-located ROIs (**Suppl. Fig. 3b-d**). More specifically, Off ACs’ local surrounds became less tuned to UV, whereas On ACs’ centre and local surround acquired a stronger preference for UV (**Suppl. Fig. 4c**). Disrupting the On pathway, on the other hand, strongly affected RF polarity of On ACs, but also strengthened the antagonistic far-surround of both On and Off ACs across the retina. The chromatic tuning of both On and Off ACs was largely unaffected (**Suppl. Fig. 3f,g,h**; **Suppl. Fig. 4d**).

While the observed changes in AC response polarity following manipulation of the On and Off pathways were expected, the difference in spatial scale was surprising: Blocking the Off pathway affected only the Off ACs’ centre response; disrupting the On pathway affected both the On ACs’ centre and local-surround responses, suggesting a substantial difference in spatial RF size and/or organisation between GABAergic On and Off ACs. The overall effects of excitatory pathway manipulations on chromatic tuning were smaller but also more diverse (cf. **Suppl. Fig. 3d,h**).

### Biologically-inspired circuit model captures chromatic responses of bipolar cells and amacrine cells

To further explore the complex interactions between BCs and ACs, we developed a biologically-inspired circuit model of the inner retina extended from Schröder et al. (2020) to study the circuits giving rise to the chromatic tuning of ACs. We focused our model on the ventral retina, thus the model included six “types” of photoreceptors (UV and green in centre, local- and far-surround), ventral BCs and all AC clusters (**Fig. 6a**) and predicted glutamate and Ca^2+^ responses of BC and AC clusters, respectively (**Fig. 6b**).

**Fig. 6.**
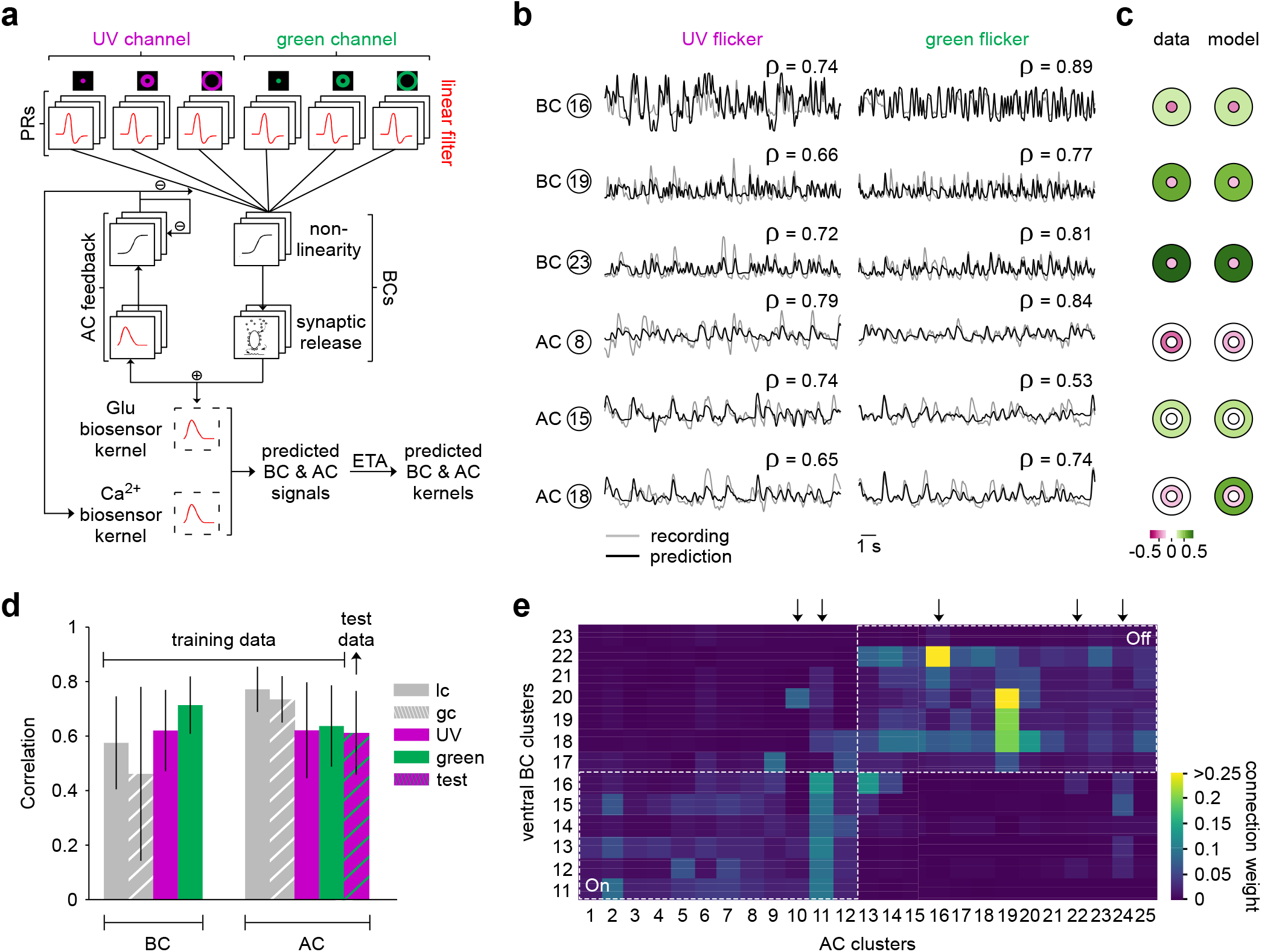
Network model of the inner retina. **(a)** Extended network model (Schröder et al., 2020) including spatio-chromatic photoreceptors (PRs), bipolar cells (BCs) and amacrine cells (ACs). Model includes ventral BC clusters and all AC clusters. BCs receive input from linear PRs and are modelled using a non-linearity and a ribbon synapse model. ACs are modelled using a linear and non-linear part, receive input from BCs, and provide inhibitory feedback to BCs and ACs. Glutamate (BC) and Ca^2+^ (AC) predictions were obtained by convolution of the model output with the iGluSnFR and GCaMP6f biosensor kernels, respectively (see Methods). Model kernels are estimated using Event Triggered Averaging (ETA) from predicted BC and AC signals. **(b)** Recording (grey) and model prediction (black) in response to UV and green flicker with Pearson correlation coefficient for representative BC and AC clusters. **(c)** Chromatic preference estimated as spectral contrast (*SC*) for data (left) and model predictions (right) of the BC and AC clusters shown in (b). **(d)** Pearson correlation coefficient (mean *±*s.d.) of BC and AC clusters for training stimuli (local chirp (lc), global chirp (gc), UV and green flicker) and AC performance for a withheld test sequence (see **Fig. 1c**) including UV and green stimulation. **(e)** Model weights of BC to AC connectivity matrix. Dashed lines separate On and Off clusters. Arrows indicate AC clusters with multi-stratified IPL profile.

For simplicity, the model did not include glycinergic ACs and we did not explicitly model horizontal cells (see Methods for details). Also, as we only modelled three spatial regions at the level of photoreceptors, the model lacked a realistic large-scale spatial structure. We optimised model parameters using a deep learning approach with the objective to maximise the correlation between model responses and recordings to achromatic (local/global chirp) and chromatic (UV/green flicker) stimuli and to reproduce the *SC* of clusters. In addition, the objective encompassed regularisation terms, including a constraint on the connectivity between clusters based on their co-stratification in the IPL to encourage biologically plausible connectivity in the model.

After training, we found that model predictions exhibited high correlation with recordings of chromatic flicker responses for both ACs and BCs and that performance assessed on AC data not seen during training was on par with training performance (**Fig. 6b,d**). In addition, the *SC* of model predictions matched the *SC* of experimental data (Pearson correlation coefficient, BC centre and far surround (0.42, 0.72); AC centre, local- and far-surround (0.7, 0.31, 0.51); **Fig. 6c**), and the BC to AC connectivity matrix exhibited a structure resembling the co-stratification profile of clusters in the IPL (**Fig. 6e**). Most On and Off AC clusters received input from On and Off BC clusters, respectively. However, some AC clusters received input from BCs of both polarities, among which were most of the AC clusters with a multi-stratified IPL profile (*C*10, *C*11, *C*16, *C*22, *C*24) – clusters with less than 70% of ROIs in On/Off layer were considered multi-stratified (indicated by arrow in **Fig. 6e**). Thus, the circuit model captured chromatic and achromatic response properties of the IPL network well.

### *In silico* pharmacological manipulations

Next, we performed *in silico* pharmacological manipulations on the circuit model to test the contribution of different circuit components to the chromatic response properties of AC clusters. We reasoned that if the model showed a similar effect as the pharmacology data to identical manipulation (e.g., disrupting the On pathway), this would provide evidence that the underlying circuit structure was well captured by the model. If this was not the case, this would suggest that circuit components not captured in our model were involved in mediating the pharmacological effect. Therefore, we predicted responses after selectively removing GABAergic AC inhibition, On BC or Off BC input from the network (**Fig. 7a** top, middle and bottom, respectively) by setting their connectivity matrices to zero. We separated clusters into On and Off using the *OOi* of the “centre-control” condition and examined how their *OOi* (**Fig. 7a,c**) and *SC* (**Fig. 7b,c**) changed following the *in silico* pharmacology and compared the effects to data from ventral ROIs from the *ex vivo* interventions described above.

**Fig. 7.**
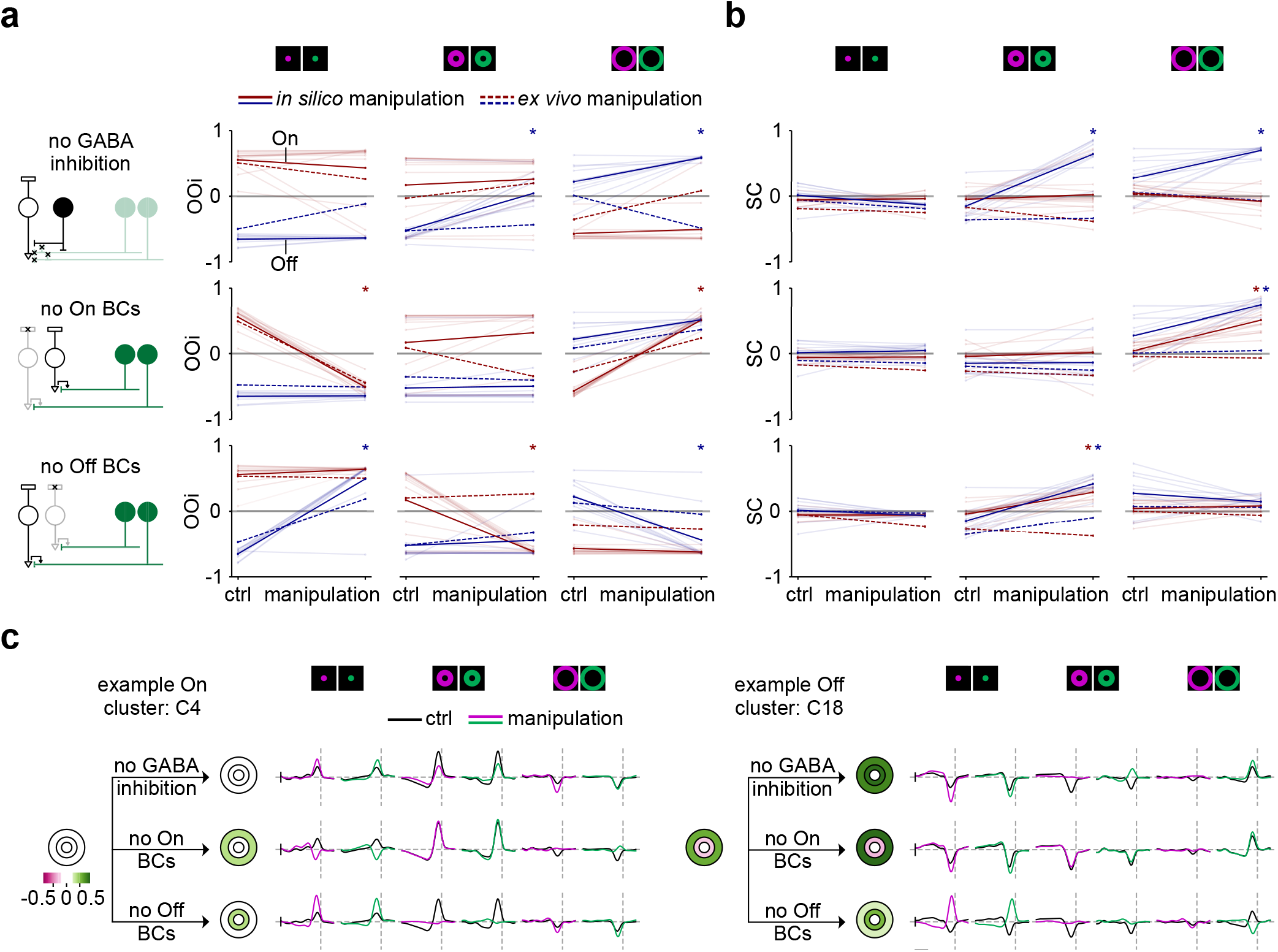
Effects of *in silico* pharmacology on amacrine cell chromatic responses. **(a)** On-Off index (*OOi*) in *in silico* control and manipulation condition in On and Off AC clusters (red and blue solid lines, respectively) from the ventral model (Fig. 6) for centre (left), local- (middle), and far-surround (right). We used the *OOi* of the “centre-control” condition to define On and Off clusters. Top: *In silico* removal of inhibition by setting AC to AC and AC to BC feedback to zero. Middle: *In silico* removal of On BCs by setting photoreceptor (PR) to On BC and On BC to AC signalling to zero. Bottom: *In silico* removal of Off BCs by setting PR to Off BC and Off BC to AC signalling to zero. Dashed lines indicate data averages from ventral ROIs for the three drug conditions (from top to bottom: application of TPMPA/gabazine, L-AP4, and UBP 310) shown in Fig. 5 and Suppl. Fig. 3). \**p <* 0.05; paired t-test corrected for multiple comparisons. **(b)** Same as in (a) but for spectral contrast (*SC*). **(c)** *SC* of *in silico* control and manipulation conditions and UV and green temporal kernels for centre, local-, and far-surround for *in silico* control (black) and manipulation (coloured) conditions for a representative On (*C*4) and Off (*C*18) AC cluster.

First, for the AC centre responses, we observed that all three *in silico* manipulations affected *OOi* and *SC* similarly to the *ex vivo* pharmacology interventions (**Fig. 7a,b,c**) with the exception of the Off ACs after removing GABA inhibition. There, the model predicted no change in response polarity, while in the retina we observed such a change. Overall, this indicates that our model likely captured essential circuit components in the centre of the simulated circuit. Next, for the local- and far-surround AC responses, we observed some similarities but also several interesting differences between model and pharmacology data. For instance, for the far-surround, the model captured the trends in *OOi* changes for both On and Off AC responses when disrupting the BC pathways, but did not predict the switch in response polarity for the no-GABA condition (**Fig. 7a right**). Most differences between model and data we found for the *SC* in the local- and far-surround. For example, upon removal of GABAergic inhibition, the *SC* of almost all modelled Off ACs increased, indicating stronger green tuning in the surround of these cells (**Fig. 7b**), in contrast to the pharmacological data, which exhibited shifts toward UV or no change. Similarly, *SC* of On and Off ACs increased in the model uniformly cross AC clusters for the far- and local-surround when blocking On and Off BCs, respectively, while it remained rather constant in the pharmacology data.

Together, this suggests that the spatial BC-AC interactions are more complex than what can currently be captured by the simple spatial layout of our model and that some of the crucial circuit components responsible for keeping ACs diversely tuned, and not simply following BC surround tuning, are not yet present in our model (see Discussion).

## Discussion

We studied the chromatic RFs of GABAergic ACs by imaging Ca^2+^ signals in their processes, which allowed us to systematically characterise their chromatic tuning properties across the retina. We found that the chromatic tuning of ACs was much more diverse than that of BCs, the ACs’ main source of excitatory inputs. By clustering the AC responses into functional types, we found 25 clusters that differed in their achromatic RF organisation (i.e. response polarity and centre vs. surround RF size), and in their chromatic tuning. In many clusters, chromatic RFs differed between the dorsal and ventral retina, suggesting that amacrine cells with similar achromatic RF organisation may play different roles in chromatic processing in dorsal vs. ventral retina. We sought to understand how the chromatic tuning of ACs is established by manipulating their synaptic inputs using pharmacology and a biologically-inspired deep learning model. Based on these data, we propose that the diverse chromatic RF properties of ACs are established through a complex interplay between the BCs’ chromatic tuning and inhibition from ACs.

### Chromatic information flow through the retina

Mouse cone photoreceptors form a gradient of opsin expression that establishes separate domains of chromatic processing in the dorsal and ventral retina. These differences in cone spectral sensitivity lead to distinct chromatic preferences in the down-stream BCs. Classifications of BCs based on different modalities agree that there are 14 BC types of in the mouse retina (Behrens et al., 2016; Franke et al., 2017; Helmstaedter et al., 2013; Kim et al., 2014; Shekhar et al., 2016). Here, by clustering BCs based on their chromatic tuning, we found twice the number of BC types, with two almost equally-sized sets of BC type segregated into dorsal and ventral based on their chromatic preferences. This confirms that most BC types simply mirror the spectral tuning of the local cone population. We observed only one BC cluster that exhibited a broad retinal distribution and stratified close to the IPL border with the GCL, which likely represents the type 9 S-cone selective BC (Behrens et al., 2016; Breuninger et al., 2011; Haverkamp et al., 2005).

Unlike BCs, ACs did not, for the most part, exhibit this segregation into dorsal and ventral groups: almost all AC clusters were broadly distributed along the dorso-ventral axis. In addition, we found that ACs displayed much weaker but also much more diverse chromatic tuning than the BCs. Our results suggest that some ACs are well-suited to remove the chromatic bias (de-bias) coming from BCs, and thereby enable chromatically more balanced RGC responses (Szatko et al., 2020). In line with our results, recent work in zebrafish retina proposed a similar role for ACs (Wang et al., 2023); they found that feedback from ACs altered the BCs’ chromatic tuning, rendering some BCs more strongly and others more weakly tuned. Together, this suggest that the need for re-balancing biased chromatic channels is not a mouse-specific function of AC circuits.

A major question we sought to address is how ACs, despite receiving their excitatory drive from BC, achieve such different chromatic RF organisation and tuning. Together, pharmacology and modelling point towards an important role of medium to long-range inhibition, some of which was not captured by our spatially-confined model. This could be either mediated by horizontal cells already in the outer plexiform layer (OPL) or by medium-field GABAergic ACs providing inhibition from outside the modelled circuit to the surround. Lateral inhibition could also be provided by polyaxonal ACs, whose axons can span over long distances across the retina (Diamond, 2017) and therefore could relay chromatic signals between dorsal and ventral retina (e.g., Chang et al., 2013). Glycinergic crossover inhibition did not seem to play a role in chromatic processing under our conditions and rather shaped achromatic properties of the ACs. Finally, a potential source of green centre responses in ventral ACs are rod photoreceptors, which provide green-sensitive input though the rod BCs across the retina.

RGCs exhibit a variety of chromatic preferences, with a distinct population of ventral RGCs showing colour opponency (Joesch and Meister, 2016; Szatko et al., 2020), which presumably support colour vision in the upper visual field (Denman et al., 2018). Recent work from our group found that the chromatic tuning of several RGC types cannot be explained simply based on their BC inputs (Szatko et al., 2020), with some RGC types displaying less colour-opponency and others more than expected. Another recent study found that several RGC types process chromatic information in a non-linear manner that depended on inhibition from the surround (Khani and Gollisch, 2021). Together, these results suggest that an RGC’s tuning is shaped by the integration of differently-tuned chromatic units. The diverse spatio-chromatic tuning we found in the ACs may well play a crucial role in shaping RGC RF properties, through enhancing or de-biasing chromatic information compared to other visual features.

### Chromatic processing by amacrine cells

We observed several tantalising motifs in the ACs’ RF features that suggest distinct roles in chromatic processing in the retina. First, we uncovered one On cluster (*C*12) and one Off cluster (*C*13) that were highly UV-sensitive and restricted to the ventral retina. In addition, there were only two clusters with UV sensitivity in the far-surround (*C*13, *C*22). These ACs could represent S-cone selective ACs, which have been described in primate (Patterson et al., 2020) and ground squirrel (Chen and Li, 2012). In addition, we observed one cluster that was green-sensitive (*C*24), although many clusters included green-sensitive ROIs in the dorsal retina. Finally, we observed two clusters that were colour-opponent (*C*18, *C*19). Both of these clusters were Off in the centre, with the local-surround responding strongly to the offset of UV light and the far-surround responding strongly to the onset of green light. This contrasts with the zebrafish, where colour-opponency in ACs was predominantly found in the On pathway (Wang et al., 2023).

In general, our experimental data suggest that there may be a spatial offset in how ACs represent UV vs. green chromatic information transmitted from BCs. In particular, green information remains local to the centre of AC RFs in the dorsal retina and UV information moves laterally into the local surround in the ventral retina (see also **Fig. 4b**). This difference in the spatial representation of chromatic signals could have important implications for spatial processing in the dorsal vs. ventral retina. Our model only included three spatial domains at the level of the PRs, while the other cells modelled are not spatially extended. As a result, we are likely not capturing the potential mechanisms underlying spatial integration with our model. Further studies using more specific circuit manipulations to study chromatic processing, such as chemogenetics, would aid in a more detailed understanding of specific cell type contributions.

Our model captures effects of pharmacological manipulations on chromatic signals in the centre better than in the local and far surround. Across clusters, the far-surround of RFs was largely achromatic, excepting the clusters mentioned at the beginning of this section. This achromatic, antagonistic surround was sometimes coupled with different chromatic preferences in the centre and local-surround, even though both centre and local-surround regions preferred the same stimulus polarity (*C*18, *C*19; **Fig. 4c**). Thus, a complex RF for colour vs. contrast emerges in many ACs, which is expected to be capable of detecting higher spatial frequencies for colour contrast than luminance contrast.

### Diversity of amacrine cell types

Cataloguing the diversity of AC types represents a major focus for the field of retinal neurobiology. Recent transcriptomic studies identified 43 (Yan et al., 2020) and 52 (Li et al., 2024) GABAergic molecular AC types. An electron microscopic survey uncovered 33 AC types (out of 45) (Helmstaedter et al., 2013) that are medium/wide field and therefore most likely also GABAergic; since they used a relatively small tissue volume, they likely missed some wide-field types. Functional classification of the ACs, however, has presented a challenge because ACs are primarily non-spiking and perform computations in their processes, and hence, cannot be easily identified by their somatic responses.

Now, multiple groups have made functional classifications of ACs similar to what has already been available for BCs (Franke et al., 2017) and RGCs (Baden et al., 2016). We used PCA and GMM clustering to provide a systematic characterisation of the chromatic signals in GABAergic ACs in the mouse retina, identifying 25 functional types. Another recent study has examined the achromatic spatio-temporal properties of mouse ACs by imaging their GABA release with a fluorescent genetically-encoded GABA sensor (Matsumoto et al., 2024) and found 44 functional types. This suggests that either ACs are more diverse in their spatio-temporal RFs than in their chromatic RFs, or that some AC types did not respond at all to our chromatic stimuli. Another notable difference between these studies is the use of different sensors, which effectively monitor different stages of neuronal signalling. GABA represents the final output of the AC computational process. Ca^2+^ signals, on the other hand, may not always be correlated with the functional output of AC processes, as Ca^2+^ can come from different sources, both extra- and intracellular, or may represent dendritic voltage or synaptic release in a nonlinear manner (Tran-Van-Minh et al., 2015). This said, Ca^2+^ signals typically correlate strongly with the GABAergic output due to the Ca^2+^ channels’ proximity and control over GABA release. Therefore, Ca^2+^ can be used as a proxy for AC output for cell types like starburst ACs (e.g., Euler et al., 2002), and we recently observed that Ca^2+^ signals recorded at the population level reveal similar direction selectivity (Strauss et al., 2022) as what was now confirmed with GABA imaging (Matsumoto et al., 2024). Further investigation of chromatic processing in ACs using a GABA sensor will be an instructive next step for understanding the relationship between dendritic Ca^2+^ signalling and GABA release.

Several themes about AC processing were observed from both Ca^2+^ and GABA imaging studies. Both studies found that using the contrast and temporally-modulated chirp stimulus was not very effective at clustering the ACs into functional groups. Rather, spatial features of the RF(chromatic or achromatic) were more important for classification. This is perhaps not surprising given the wide variety of sizes and morphologies of AC dendritic arbours and their long-described role in lateral inhibition (Diamond, 2017). In both studies, we found a wide variety of complex RFs. Matstumoto and coworkers (Matsumoto et al., 2024) note that this wide variety allows ACs to sample and encode a large range of spatial and temporal frequencies. Finally, in both studies, some functional clusters spanned both On and Off layers, suggesting that there are GABAergic ACs that perform “crossover inhibition”, a function classically attributed to glycinergic ACs (Molnar and Werblin, 2007). In general, how our classification overlays with the classification based on GABA release, will be important for understanding the specific functions of ACs. In particular, how the 40+ spatiotemporal types overlay with the 25 chromatic types described here will be an important next step in understanding the functional roles of the retina’s most diverse cell class, may also contribute to a better understanding the roles of interneurons in other parts of the brain.

## Methods

### Animals and tissue preparation

All animal procedures were approved by the governmental review board (Regierungspräsidium Tübingen, Baden-Württemberg, Konrad-Adenauer-Str. 20, 72072 Tübingen, Germany) and performed according to the laws governing animal experimentation issued by the German Government. For Ca^2+^ imaging in the IPL, we used the transgenic line STOCK Gad2tm2(cre)Zjh/J (Gad2-IRES-Cre, JAX 010802, The Jackson Laboratory; see Taniguchi et al., 2011) that was crossbred with the Cre-dependent green fluorescent reporter line B6;129S-Gt(ROSA)26Sortm95.1(CAG-GCaMP6f)Hze/J (Ai95D, JAX 024105; see Madisen et al., 2015), which expresses GCaMP6f. We used mice between 5 and 16 weeks old of either sex (*n* = 17 mice). Owing to the exploratory nature of our study, we did not use randomisation and blinding. No statistical methods were used to predetermine sample size.

Animals were housed under a standard 12 h day/night rhythm at 22° C and 55% humidity. For our recordings, the animals were first dark-adapted for at least 1 h, then anaesthetised with isoflurane (Baxter) and sacrificed by cervical dislocation. All procedures described further are performed under very dim red light (>650 nm).

The eyes were removed and the retina was extracted in carboxygenated (95% O_2_, 5% CO_2_) artificial cerebrospinal fluid (ACSF) solution (containing in mM: 125 NaCl, 2.5 KCl, 2 CaCl_2_, 1 MgCl_2_, 1.25 NaH_2_PO_4_, 26 NaHCO_3_, 20 glucose, and 0.5 L-glutamine (pH 7.4)). The retina was then transferred to the recording chamber, which was constantly perfused with carboxygenated, *≈*36° C warm ACSF containing 0.1 μM Sulforhodamine-101 (SR101, Invitrogen) to reveal the retina’s vasculature and any damaged cells in the red fluorescent channel (Euler et al., 2009). The dorso-ventral axis of the retina was marked to trace the position of the recorded fields relative to the optic disc.

### Two-photon imaging

We used a MOM-type two-photon microscope (designed by W. Denk, MPI, Heidelberg; purchased from Sutter Instruments/Science Products, see Euler et al., 2009). In brief, the system was equipped with a mode-locked Ti:Sapphire laser tuned to 927 nm (MaiTai-HP DeepSee, Newport Spectra-Physics), two fluorescence detection channels for GCaMP6f (HQ 510/84, AHF/Chroma) and SR101 (HQ 630/60, AHF), and a water immersion objective (W Plan-Apochromat ×20 /1.0 DIC M27, Zeiss). For vertical image acquisition, we used custom-made software (ScanM by M. Müller and T.E.) running under IGOR Pro 6.37 for Windows (Wavemetrics) and an electrically tunable lens (Zhao et al., 2020) recording time-lapsed 64 × 56 pixel image scans (at 11.16 Hz).

### Light stimulation

A DLP projector (lightcrafter, DPM-E4500UVBGMKII, EKB Technologies Ltd) with internal UV and green light-emitting diodes (LEDs) was focused through the objective. The LEDs were band-pass filtered (390/576 Dualband, F59-003, AHF/Chroma), for achieving an optimal spectral separation of mouse M- and S-opsins, and were synchronised with the microscope’s scan retrace. Photoisomerisation rates were set to range from *≈*0.5 (black image) to *≈*20 x 10^3^ P* per s per cone for M- and S-opsins, respectively (for details, see Franke et al., 2019). In addition, a steady illumination component of 10^4^ P* per s per cone was present during the recordings because of two-photon excitation of photopigments (discussed in Baden et al., 2013; Euler et al., 2019, 2009). To allow the retina to adapt to the laser, the tissue was scanned for at least 15 s before the stimulus was presented. The stimulus was centred to the recorded field.

Three types of light stimuli were used: (a) a chromatic flicker stimulus consisting of three concentric regions: *centre* (100 μm in diameter), *local-surround* (a 300 μm annulus sparing the central 100 μm), *far-surround* (a 800 μm annulus sparing the central 300 μm) that flickering independently for UV or green in a random binary sequence at 5 Hz (modified from Szatko et al., 2020). Two minute epochs of either UVor green stimulation were interleaved with repeated test sequences of 7 s green and 7 s UV. The test sequences were used to validate the model and were not used in calculating the receptive field kernels; (b) local chirp (100 μm in diameter; see Franke et al., 2017); and (c) full-field chirp stimulus (800 μm in diameter; see Baden et al., 2016).

### Pharmacology

For the pharmacological experiments, we used: 75 μM TPMPA ((1,2,5,6-Tetrahydropyridin-4-yl)methylphosphinic acid); 10 μM gabazine (SR-95531); 0.5 μM strychnine; 10 μM UBP 310 ((S)-1-(2-Amino-b 2-carboxyethyl)-3-(2-carboxy-thiophene-3-yl-methyl)-5-methylpyrimidine-2,4-dione); and 50 μM L-AP4 (L-2-amino-4-phosphonobutyric acid). The drug solutions were carboxygenated before application. Control recordings were made, and then drugs were bath-applied for 15 min before recording in drug conditions.

### Data analysis

Data were pre-processed using IGOR Pro (WaveMetrics, v8.04), organised and further analysed in Python 3 and a custom-written database schema in Data-joint (Yatsenko et al., 2015). We defined ROIs using custom correlation-based algorithms. ROI sizes were restricted to range from 0.8 to 4 μm in diameter. Correlation thresholds were determined separately for each scan line, to correct for variability of GCaMP6f labelling and laser intensity across the IPL (Zhao et al., 2020). For each field-of-view, we manually defined the IPL borders with the INL (= 0) and the GCL (= 1). Then, each ROI’s location in the IPL was calculated as the distance of the ROI centre to the IPL borders. ROIs outside the IPL borders were excluded.

The Ca^2+^ traces for each ROI were extracted (as Δ*F/F*), detrended with a high-pass filter of 0.2 Hz and znormalised by subtracting each traces’ mean and dividing by its s.d. Stimulus time markers embedded in the recorded data served to align each ROI’s trace to the visual stimulus with 1.6-ms precision. For this, the timing for each ROI relative to the stimulus was corrected for sub-frame time-offsets related to the scanning. We used linear interpolation to resample response traces to line precision (625 Hz). Responses to repeated trials of the local and global chirp stimulus were binned at 10 Hz and averaged across measurements within a bin.

#### Temporal kernels

To estimate temporal kernels, we first z-scored the up-sampled response traces and stimulus traces of the UV and green flicker. For each condition, we calculated the dot product of response trace and stimulus trace and normalised it by the number of data points. We repeated this procedure using time-shifted stimulus traces to obtain 2-s temporal kernels. We excluded time periods of the test sequence for kernel estimation.

#### Stimulus Artefact

During our analysis, we observed an artefact in the UV far-surround temporal kernels of some ROIs, manifesting as a peak in the signal prior to the response time. This artefact was likely due to UV light exciting the GCaMP fluorophore directly. To remove this artefact, we first estimated UV far-surround temporal kernels for the 5% least-responding single pixels (based on the s.d. of the response) from five randomly picked scan fields. We averaged them to obtain an “artefact temporal kernel”. Next, we convolved this kernel with the UV far-surround stimulus and subtracted the resulting artefact-related response component weighted by a ROI-specific value *α* from the ROIs’ response traces to obtain a corrected trace for each ROI. We estimated *α* for each ROI such that it would minimise

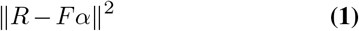

where *R* is the ROI’s response and *F* the artefact response, yielding the following solution We set negative values of *α* to zero and finally, re-estimated each ROI’s temporal kernels using the corrected traces.

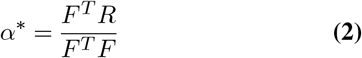

#### Response quality

To estimate kernel quality, we measured the kernel amplitude as the difference between the maximum and minimum response values in a time window of 700 ms relative to the response time. ROIs with at least one kernel (UV or green centre, UV or green local-surround, green far-surround) exceeding kernel quality of 0.1 are used for further analysis (*n* = 5, 378 of 6, 628). Note that the UV far-surround kernel was excluded from quality filtering because of the aforementioned artefact. For analysis of the paired-data from the drug experiments, we included ROIs that exceeded the centre kernel quality threshold of 0.1 in one of the four cases: UV/green, control/drug (TPMPA/gbz: *n* = 546*/*709, strychnine: *n* = 497*/*586, L-AP4: *n* = 200*/*269, UBP 310: *n* = 445*/*513).

#### Spectral contrast

To estimate chromatic preference of ROIs, we calculated a spectral contrast (*SC*) for each spatial condition (centre, local-surround, far-surround) using kernel amplitudes (*A*) for UV and green. *SC* was calculated as Michelson contrast as follows:

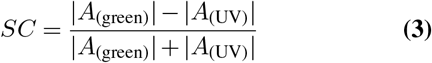

#### On-Off index

To estimate response polarity of ROIs, we calculated the On-Off index (*OOi*) for each spatial condition (centre, local-surround, far-surround). First, each kernel was convolved with a stimulus consisting of 1 s On and 1 s Off steps and the average response between UV and green was calculated. Next, the mean achromatic response (*r*) for the On and the Off window was calculated. *OOi* was calculated as Michelson contrast as follows:

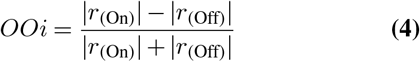

#### Hierarchical clustering

To investigate relationships between chromatic AC clusters, we normalised average clusters’ response traces for all chromatic stimuli by dividing by the maximum value across stimuli, calculated the Mean Squared Error between cluster pairs, and performed hierarchical clustering (`scipy.cluster.hierarchy.linkage, method= ‘weighted’`, see Virtanen et al., 2020).

#### BC Dataset

The recording and processing of the BC dataset (Szatko et al., 2020) was almost identical to that of the AC dataset, with the following differences: The BC chromatic flicker consisted of only two concentric regions: *centre* (100 μm in diameter) and *far surround* (a 800 μm annulus sparing the central 100 μm), flickering at 10 Hz, and it did not contain test sequences. The duration of the UV and green flicker presented to BCs was 180 s each. We did not exclude ROIs outside the IPL borders as BCs were imaged with a glutamate sensor and, thus, signals were related to extracellular glutamate and not specific neuronal structures. Also, the UV far-surround temporal kernels did not contain a stimulus artefact. We estimated the kernel quality in a time window 380 ms relative to the response time and all four conditions (UV and green, centre and far-surround) were used to determine kernel quality. All data was reprocessed for the present study to ensure consistent pre-processing.

### Dimensionality reduction and clustering

To identify functional cell types, we followed Baden et al. (2016). Before clustering, we extracted features using Principal Component Analysis (PCA; `sklearn.decomposition.PCA`) on the six temporal RFs (UV and green centre, local- and far-surround) independently. We included the time period before the response time and examined the explained variance as a function of the number of components (between 1 and 6 components). We determined that the “elbow” (Cattell, 1966) occurred at 2 or 3 components for each of the six conditions and, therefore, continued our analysis with 2 and 3 components in parallel.

Next, we z-scored the features we obtained with PCA and clustered ROIs using a Gaussian Mixture Model (GMM; `sklearn.mixture.GaussianMixture`). We carried out the clustering with four different covariance structures (*full, spherical, diagonal*, and *tied*) and tested between 1 and 50 components. In addition, we ran each of those 200 configurations with 20 different random initialisations. We calculated the Bayesian Information Criterion (BIC) for each model and averaged across the random initialisations to obtain an average BIC curve for each covariance type. Next, we calculated the Bayes Factor as *BF* = 2 *·* Δ_*BIC*_ between sub-sequent cluster numbers. We stopped increasing the number of clusters when BF was smaller than 6 (Baden et al., 2016) and determined the number of clusters for each average BIC curve in this manner. We then took the clustering with the smallest BIC from the 20 random initialisations with the specific number of clusters determined from the *BF*.

To choose between the different covariance types and number of PCA features, we visually examined the resulting clusters. We excluded the models with a full covariance matrix due to their small number of clusters (6 and 5 clusters for the models with 2 and 3 PCA features, respectively), and the ones with a spherical covariance matrix due to their high number of clusters (46 and 38 clusters for the models with 2 and 3 PCA features, respectively). Next, we excluded the models with a spherical covariance matrix, as their cluster sizes were less homogeneous compared to the models with a diagonal covariance matrix. Finally, we selected the diagonal model which was trained with 3 PCA features due to its number of clusters (25 clusters compared to 35 for the model trained using 2 PCA features). To assess the quality of our clustering, we generated 10,000 samples from the trained GMM, predicted labels for them and quantified the accuracy *AC* of those predictions for each cluster individually as

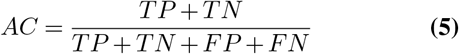

where *TP* and *TN* are True Positives and True Negatives, respectively and *FP* and *FN* are False Positives and False Negatives, respectively. We observed high accuracy (>0.97) for each of the 25 clusters, indicating that they can be confidently separated in feature space.

For the BC dataset, we excluded the models with a diagonal, tied and spherical covariance matrix due to the large number of clusters (50 for all models) and selected the model with a full covariance matrix trained with 3 PCA features due the lower number of clusters (24 compared to 30 for the model trained with 2 PCA features). Upon sampling from the trained GMM, we achieved high accuracy for all 24 clusters (>0.99).

#### Preprocessing for model training

We obtained cluster response traces for model training by averaging across responses of ROIs in a cluster for the different stimuli (local and global chirp, UV and green flicker, test sequence). For the test sequence, we additionally averaged across the 4 repetitions of the sequence. We only used cluster responses to the test sequences for model evaluation after training. For the UV and green flicker, there was one field, for which the stimulus had lower than average correlation with the stimuli from other fields. Therefore, we excluded ROIs from this field when calculating the cluster response trace for the UV and green flicker stimulus. Next, we used linear interpolation to resample cluster response traces and stimulus traces to 64 Hz. Finally, we normalised the cluster response traces for all stimuli by subtracting their mean and dividing by the maximum value across stimuli.

### Model

We modified and extended the biologically-inspired circuit model published in Schröder et al. (2020). The model included six PR populations with different chromatic properties (UV and green) and different spatial extent (centre, local- and far-surround), which were modelled as linear filters. The “PRs” summarised all processing upstream of the BC terminal, including any dendritic processing in BCs. BCs received a weighted sum of the PR output as their input and every BC received input from its own set of six PRs. The polarity of the centre PRs matched that of the BCs and the local- and far-surround PRs were of opposite polarity to model the centre-surround antagonism of BC RFs.

BCs were modelled as a non-linearity followed by a deterministic version of a ribbon synapse model (Schröder et al., 2019). In this ribbon synapse model, vesicles were released from the readily-releasable pool (RRP), which was replenished with vesicles from the intermediate pool (IP), and which in turn was replenished from the cytoplasm. ACs received input from BCs and were modelled using a double-exponential kernel, followed by a non-linearity. ACs provided inhibition to BCs and other ACs.

As the final step, the modelled BC and AC output signals were convolved with the glutamate (iGluSnFR; Marvin et al., 2013) and Ca^2+^ (GCaMP6f; Chen et al., 2013) biosensor kernels to obtain predictions of glutamate and Ca^2+^ responses in BCs and ACs, respectively. We included all 25 AC clusters and the 13 ventral BC clusters. The model was implemented in PyTorch (Paszke et al., 2019) and had a total of 1,637 free parameters, which included the four connectivity matrices (PR-to-BC, BC-to-AC, AC-to-BC, and AC-to-AC), a stimulus bias and scaling parameter, as well as a speed parameters for the PR kernels. In addition, we learned the parameters of the non-linearities (offset and slope) and the rise and decay time constants of the AC kernels. In the ribbon synapse model, the steady state fraction and maximum capacity of the RRP and IP, as well as their replenishing rates were learned during training.

#### Model loss

During training, we predicted model responses to all stimuli (local and global chirp, AC UV and green flicker, and BC UV and green flicker), and matched the duration of the generated model data to the duration of the experimentally recorded data. We only predicted local and global chirp responses once, while the *ex vivo* experiments included repeated trials of chirp stimuli. We then calculated the correlation between model prediction and experimental recording for all AC and BC clusters and stimuli and estimated temporal RFs and *SC* from the model flicker responses.

The training loss consisted of the multiple terms. The first term in the loss function was the mean correlation for local and global chirp, and AC and BC UV and green flicker – specifically, [1 – mean correlation], as we minimised the loss. Moreover, the loss included the MSE between experimentally recorded *SC* and model *SC* for ACs and BCs. Lastly, we included regularisation terms of the s.d. of PR kernel speeds, s.d. of BC and AC release means (before convolution with biosensor kernels), and s.d. and weights of the four connectivity matrices. We scaled the weights of the BC-to-AC, AC-to-BC, and AC-to-AC connectivity matrices while calculating the loss according to the co-stratification of cluster pairs in the IPL obtained from the experimental data to enforce biologically plausible connections in the model. We also included weights on the *SC* mean squared error and regularisation terms in the loss as hyper-parameters.

#### Model training

We optimised model parameters using the Adam optimiser (Kingma and Ba, 2017), trained a number of models with different hyper-parameters, and chose the model with the highest performance on the test set. We used the following training schedule to optimise model parameters. We began training with an initial learning rate *λ* and once the loss did not improve for *n* steps, we lowered the learning rate by 0.5 and continued training. We stopped training after lowering the learning rate *m* times or a maximum number of *t* steps had been reached. The model we chose with the best performance was trained with an initial learning rate *λ* = 0.8, *n* = 3, *m* = 8, and *t* = 400, and converged at around 200 steps.

#### Pharmacological interventions

We performed three *in silico* pharmacological interventions: (1) Removing GABAergic AC inhibition by setting AC-to-BC and AC-to-AC connectivity matrices to zero; (2) Removing On BCs by setting PR-to-BC and BC-to-AC matrices for On BCs to zero: the polarity of BCs was determined by their centre temporal RF; (3) Removing Off BCs by setting PR-to-BC and BC-to-AC matrices for Off BCs to zero.

### Statistical analysis

We used paired t-test corrected for multiple comparisons to quantify the difference between control and drug condition. We did so for *SC* and *OOi* in different spatial and retina locations and split ROIs into On and Off (**Fig. 5**; **Suppl. Fig. 3**; **Suppl. Fig. 4**).

## Acknowledgements

We thank Gordon Eske and Merle Harrer for excellent technical assistance. Funded by the German Research Foundation (DFG; BE 5601/2-1; SPP 2041; BE 5601/4-1,2; EU 42/9-1,2; EU 42/12-1), the German Ministry of Education and Research (BMBF; Bernstein Award 01GQ1601; BCCN 01GQ1002), the Medical Faculty/U Tübingen (fortüne), and the Max Planck Society (MPG; M.FE.A.KYBE0004).

## Author Contributions

Conceptualisation: MK, SS, AV, TE, KF, PB; Data curation: MK, SS; Formal analysis: MK, SS, AV; Investigation: MK, SS; Methodology: MK, SS; Funding acquisition: AV, TE, PB; Project administration: TS, TE, PB; Resources: TE, PB; Software: MK, SS, AV, KF; Supervision: AV, KF, TE, PB; Validation: MK, SS; Visualisation: MK, SS; Writing – original draft: MK, SS, AV; Writing – review and editing: all authors.

## References

Applebury, M., Antoch, M., Baxter, L., Chun, L., Falk, J., Farhangfar, F., Kage, K., Krzystolik, M., Lyass, L., and Robbins, J. (2000). The Murine Cone Photoreceptor. Neuron, 27(3):513–523.

Baden, T., Berens, P., Franke, K., Román Rosón, M., Bethge, M., and Euler, T. (2016). The functional diversity of retinal ganglion cells in the mouse. Nature, 529(7586):345–350. Number: 7586 Publisher: Nature Publishing Group.

Baden, T., Nikolaev, A., Esposti, F., Dreosti, E., Odermatt, B., and Lagnado, L. (2014). A synaptic mechanism for temporal filtering of visual signals. PLoS Biology, 12(10):e1001972.

Baden, T. and Osorio, D. (2019). The Retinal Basis of Vertebrate Color Vision. Annual Review of Vision Science, 5(1):177–200.

Baden, T., Schubert, T., Berens, P., and Euler, T. (2018). The Functional Organization of Vertebrate Retinal Circuits for Vision. In Oxford Research Encyclopedia of Neuroscience. Oxford University Press.

Baden, T., Schubert, T., Chang, L., Wei, T., Zaichuk, M., Wissinger, B., and Euler, T. (2013). A Tale of Two Retinal Domains: Near-Optimal Sampling of Achromatic Contrasts in Natural Scenes through Asymmetric Photoreceptor Distribution. Neuron, 80(5):1206–1217.

Behrens, C., Schubert, T., Haverkamp, S., Euler, T., and Berens, P. (2016). Connectivity map of bipolar cells and photoreceptors in the mouse retina. eLife, 5:e20041.

Borghuis, B. G., Looger, L. L., Tomita, S., and Demb, J. B. (2014). Kainate Receptors Mediate Signaling in Both Transient and Sustained OFF Bipolar Cell Pathways in Mouse Retina. The Journal of Neuroscience, 34(18):6128–6139.

Breuninger, T., Puller, C., Haverkamp, S., and Euler, T. (2011). Chromatic Bipolar Cell Pathways in the Mouse Retina. The Journal of Neuroscience, 31(17):6504–6517.

Cattell, R. B. (1966). The Scree Test For The Number Of Factors. Multivariate Behavioral Research, 1(2):245–276.

Chang, L., Breuninger, T., and Euler, T. (2013). Chromatic Coding from Cone-type Unselective Circuits in the Mouse Retina. Neuron, 77(3):559–571.

Chen, S. and Li, W. (2012). A color-coding amacrine cell may provide a blue-Off signal in a mammalian retina. Nature Neuroscience, 15(7):954–956.

Chen, T.-W., Wardill, T. J., Sun, Y., Pulver, S. R., Renninger, S. L., Baohan, A., Schreiter, E. R., Kerr, R. A., Orger, M. B., Jayaraman, V., Looger, L. L., Svoboda, K., and Kim, D. S. (2013). Ultrasensitive fluorescent proteins for imaging neuronal activity. Nature, 499(7458):295–300. Publisher: Nature Publishing Group.

Denman, D. J., Luviano, J. A., Ollerenshaw, D. R., Cross, S., Williams, D., Buice, M. A., Olsen, S. R., and Reid, R. C. (2018). Mouse color and wavelength-specific luminance contrast sensitivity are non-uniform across visual space. eLife, 7:e31209.

Diamond, J. S. (2017). Inhibitory Interneurons in the Retina: Types, Circuitry, and Function. Annual Review of Vision Science, 3(1):1–24.

Ekesten, B. and Gouras, P. (2005). Cone and rod inputs to murine retinal ganglion cells: Evidence of cone opsin specific channels. Visual Neuroscience, 22(6):893–903.

Euler, T. and Denk, W. (2001). Dendritic processing. Curr. Opin. Neurobiol., 11(4):415–422.

Euler, T., Detwiler, P. B., and Denk, W. (2002). Directionally selective calcium signals in dendrites of starburst amacrine cells. Nature, 418(6900):845–852.

Euler, T., Franke, K., and Baden, T. (2019). Studying a Light Sensor with Light: Multiphoton Imaging in the Retina. In Hartveit, E., editor, Multiphoton Microscopy, volume 148, pages 225–250. Springer New York, New York, NY. Series Title: Neuromethods.

Euler, T., Hausselt, S. E., Margolis, D. J., Breuninger, T., Castell, X., Detwiler, P. B., and Denk, W. (2009). Eyecup scope—optical recordings of light stimulus-evoked fluorescence signals in the retina. Pflügers Archiv - European Journal of Physiology, 457(6):1393–1414.

Feord, R. C., Gomoliszewska, A., Pienaar, A., Mouland, J. W., and Brown, T. M. (2023). Colour opponency is widespread across the mouse subcortical visual system and differentially targets GABAergic and non-GABAergic neurons. Scientific Reports, 13(1):9313.

Franke, K., Berens, P., Schubert, T., Bethge, M., Euler, T., and Baden, T. (2017). Inhibition decorrelates visual feature representations in the inner retina. Nature, 542(7642):439–444.

Franke, K., Cai, C., Ponder, K., Fu, J., Sokoloski, S., Berens, P., and Tolias, A. S. (2023). Asymmetric distribution of color-opponent response types across mouse visual cortex supports superior color vision in the sky. preprint, elife.

Franke, K., Maia Chagas, A., Zhao, Z., Zimmermann, M. J., Bartel, P., Qiu, Y., Szatko, K. P., Baden, T., and Euler, T. (2019). An arbitrary-spectrum spatial visual stimulator for vision research. eLife, 8:e48779.

Gerl, E. J. and Morris, M. R. (2008). The Causes and Consequences of Color Vision. Evolution: Education and Outreach, 1(4):476–486.

Haverkamp, S., Wässle, H., Duebel, J., Kuner, T., Augustine, G. J., Feng, G., and Euler, T. (2005). The Primordial, Blue-Cone Color System of the Mouse Retina. The Journal of Neuroscience, 25(22):5438–5445.

Helmstaedter, M., Briggman, K. L., Turaga, S. C., Jain, V., Seung, H. S., and Denk, W. (2013). Connectomic reconstruction of the inner plexiform layer in the mouse retina. Nature, 500(7461):168–174.

Jacobs, G. H., Williams, G. A., and Fenwick, J. A. (2004). Influence of cone pigment coexpression on spectral sensitivity and color vision in the mouse. Vision Research, 44(14):1615–1622.

Joesch, M. and Meister, M. (2016). A neuronal circuit for colour vision based on rod–cone opponency. Nature, 532(7598):236–239.

Khani, M. H. and Gollisch, T. (2021). Linear and nonlinear chromatic integration in the mouse retina. Nature Communications, 12(1):1900.

Kim, J. S., Greene, M. J., Zlateski, A., Lee, K., Richardson, M., Turaga, S. C., Purcaro, M., Balkam, M., Robinson, A., Behabadi, B. F., Campos, M., Denk, W., and Seung, H. S. (2014). Space–time wiring specificity supports direction selectivity in the retina. Nature, 509(7500):331–336. Publisher: Nature Publishing Group.

Kingma, D. P. and Ba, J. (2017). Adam: A Method for Stochastic Optimization. arXiv:1412.6980 [cs].

Li, J., Choi, J., Cheng, X., Ma, J., Pema, S., Sanes, J. R., Mardon, G., Frankfort, B. J., Tran, N. M., Li, Y., and Chen, R. (2024). Comprehensive single-cell atlas of the mouse retina. Pages: 2024.01.24.577060 Section: New Results.

Lin, B. and Masland, R. H. (2006). Populations of wide-field amacrine cells in the mouse retina. Journal of Comparative Neurology, 499(5):797–809.

Madisen, L., Garner, A., Shimaoka, D., Chuong, A., Klapoetke, N., Li, L., van der Bourg, A., Niino, Y., Egolf, L., Monetti, C., Gu, H., Mills, M., Cheng, A., Tasic, B., Nguyen, T., Sunkin, S., Benucci, A., Nagy, A., Miyawaki, A., Helmchen, F., Empson, R., Knöpfel, T., Boyden, E., Reid, R., Carandini, M., and Zeng, H. (2015). Transgenic Mice for Intersectional Targeting of Neural Sensors and Effectors with High Specificity and Performance. Neuron, 85(5):942–958.

Martersteck, E. M., Hirokawa, K. E., Evarts, M., Bernard, A., Duan, X., Li, Y., Ng, L., Oh, S. W., Ouellette, B., Royall, J. J., Stoecklin, M., Wang, Q., Zeng, H., Sanes, J. R., and Harris, J. A. (2017). Diverse Central Projection Patterns of Retinal Ganglion Cells. Cell Reports, 18(8):2058–2072.

Marvin, J. S., Borghuis, B. G., Tian, L., Cichon, J., Harnett, M. T., Akerboom, J., Gordus, A., Renninger, S. L., Chen, T.-W., Bargmann, C. I., Orger, M. B., Schreiter, E. R., Demb, J. B., Gan, W.-B., Hires, S. A., and Looger, L. L. (2013). An optimized fluorescent probe for visualizing glutamate neurotransmission. Nature Methods, 10(2):162–170. Publisher: Nature Publishing Group.

Masland, R. H. (2012). The tasks of amacrine cells. Visual Neuroscience, 29(1):3–9.

Matsumoto, A., Morris, J., Looger, L. L., and Yonehara, K. (2024). Diverse GABA signaling in the inner retina enables spatiotemporal coding. Pages: 2024.01.09.574952 Section: New Results.

Mills, S. L., Tian, L.-M., Hoshi, H., Whitaker, C. M., and Massey, S. C. (2014). Three Distinct Blue-Green Color Pathways in a Mammalian Retina. The Journal of Neuroscience, 34(5):1760–1768.

Molnar, A. and Werblin, F. (2007). Inhibitory Feedback Shapes Bipolar Cell Responses in the Rabbit Retina. Journal of Neurophysiology, 98(6):3423–3435.

Mouland, J. W., Pienaar, A., Williams, C., Watson, A. J., Lucas, R. J., and Brown, T. M. (2021). Extensive cone-dependent spectral opponency within a discrete zone of the lateral geniculate nucleus supporting mouse color vision. Current Biology, 31(15):3391–3400.e4.

Nadal-Nicolás, F. M., Kunze, V. P., Ball, J. M., Peng, B. T., Krishnan, A., Zhou, G., Dong, L., and Li, W. (2020). True S-cones are concentrated in the ventral mouse retina and wired for color detection in the upper visual field. eLife, 9:e56840.

Nakajima, Y., Iwakabe, H., Akazawa, C., Nawa, H., Shigemoto, R., Mizuno, N., and Nakanishi, S. (1993). Molecular characterization of a novel retinal metabotropic glutamate receptor mGluR6 with a high agonist selectivity for L-2-amino-4-phosphonobutyrate. The Journal of Biological Chemistry, 268(16):11868–11873.

Paszke, A., Gross, S., Massa, F., Lerer, A., Bradbury, J., Chanan, G., Killeen, T., Lin, Z., Gimelshein, N., Antiga, L., Desmaison, A., Köpf, A., Yang, E., DeVito, Z., Raison, M., Tejani, A., Chilamkurthy, S., Steiner, B., Fang, L., Bai, J., and Chintala, S. (2019). PyTorch: An Imperative Style, High-Performance Deep Learning Library. arXiv:1912.01703 [cs, stat].

Patterson, S. S., Kuchenbecker, J. A., Anderson, J. R., Neitz, M., and Neitz, J. (2020). A Color Vision Circuit for Non-Image-Forming Vision in the Primate Retina. Current Biology, 30(7):1269–1274.e2.

Puller, C., Ivanova, E., Euler, T., Haverkamp, S., and Schubert, T. (2013). OFF bipolar cells express distinct types of dendritic glutamate receptors in the mouse retina. Neuroscience, 243:136–148.

Pérez De Sevilla Müller, L., Shelley, J., and Weiler, R. (2007). Displaced amacrine cells of the mouse retina. Journal of Comparative Neurology, 505(2):177–189.

Rhim, I. and Nauhaus, I. (2023). Joint representations of color and form in mouse visual cortex described by random pooling from rods and cones. Journal of Neurophysiology, 129(3):619–634.

Rosa, J. M., Morrie, R. D., Baertsch, H. C., and Feller, M. B. (2016). Contributions of Rod and Cone Pathways to Retinal Direction Selectivity Through Development. Journal of Neuroscience, 36(37):9683–9695. Publisher: Society for Neuroscience Section: Articles.

Röhlich, P., Van Veen, T., and Szél, A. (1994). Two different visual pigments in one retinal cone cell. Neuron, 13(5):1159–1166.

Schnapf, J. L., Nunn, B. J., Meister, M., and Baylor, D. A. (1990). Visual transduction in cones of the monkey Macaca fascicularis. The Journal of Physiology, 427:681–713.

Schröder, C., James, B., Lagnado, L., and Berens, P. (2019). Approximate Bayesian Inference for a Mechanistic Model of Vesicle Release at a Ribbon Synapse. In Advances in Neural Information Processing Systems, volume 32. Curran Associates, Inc.

Schröder, C., Klindt, D., Strauss, S., Franke, K., Bethge, M., Euler, T., and Berens, P. (2020). System Identification with Biophysical Constraints: A Circuit Model of the Inner Retina. In Advances in Neural Information Processing Systems, volume 33, pages 15439–15450. Curran Associates, Inc.

Shekhar, K., Lapan, S. W., Whitney, I. E., Tran, N. M., Macosko, E. Z., Kowalczyk, M., Adiconis, X., Levin, J. Z., Nemesh, J., Goldman, M., McCarroll, S. A., Cepko, C. L., Regev, A., and Sanes, J. R. (2016). Comprehensive Classification of Retinal Bipolar Neurons by Single-Cell Transcriptomics. Cell, 166(5):1308–1323.e30.

Sher, A. and DeVries, S. H. (2012). A non-canonical pathway for mammalian blue-green color vision. Nature Neuroscience, 15(7):952–953.

Sonoda, T., Li, J. Y., Hayes, N. W., Chan, J. C., Okabe, Y., Belin, S., Nawabi, H., and Schmidt, T. M. (2020). A noncanonical inhibitory circuit dampens behavioral sensitivity to light. Science, 368(6490):527–531.

Stabio, M. E., Sabbah, S., Quattrochi, L. E., Ilardi, M. C., Fogerson, P. M., Leyrer, M. L., Kim, M. T., Kim, I., Schiel, M., Renna, J. M., Briggman, K. L., and Berson, D. M. (2018). The M5 Cell: A Color-Opponent Intrinsically Photosensitive Retinal Ganglion Cell. Neuron, 97(1):150–163.e4.

Strauss, S., Korympidou, M. M., Ran, Y., Franke, K., Schubert, T., Baden, T., Berens, P., Euler, T., and Vlasits, A. L. (2022). Center-surround interactions underlie bipolar cell motion sensitivity in the mouse retina. Nature Communications, 13(1).

Szatko, K. P., Korympidou, M. M., Ran, Y., Berens, P., Dalkara, D., Schubert, T., Euler, T., and Franke, K. (2020). Neural circuits in the mouse retina support color vision in the upper visual field. Nature Communications, 11(1):3481.

Taniguchi, H., He, M., Wu, P., Kim, S., Paik, R., Sugino, K., Kvitsani, D., Fu, Y., Lu, J., Lin, Y., Miyoshi, G., Shima, Y., Fishell, G., Nelson, S., and Huang, Z. (2011). A Resource of Cre Driver Lines for Genetic Targeting of GABAergic Neurons in Cerebral Cortex. Neuron, 71(6):995–1013.

Thoreson, W. B. and Dacey, D. M. (2019). Diverse Cell Types, Circuits, and Mechanisms for Color Vision in the Vertebrate Retina. Physiological Reviews, 99(3):1527–1573.

Tran-Van-Minh, A., Cazé, R. D., Abrahamsson, T., Cathala, L., Gutkin, B. S., and DiGregorio, D. A. (2015). Contribution of sublinear and supralinear dendritic integration to neuronal computations. Frontiers in Cellular Neuroscience, 9.

Virtanen, P., Gommers, R., Oliphant, T. E., Haberland, M., Reddy, T., Cournapeau, D., Burovski, E., Peterson, P., Weckesser, W., Bright, J., van der Walt, S. J., Brett, M., Wilson, J., Millman, K. J., Mayorov, N., Nelson, A. R. J., Jones, E., Kern, R., Larson, E., Carey, C. J., Polat, I., Feng, Y., Moore, E. W., VanderPlas, J., Laxalde, D., Perktold, J., Cimrman, R., Henriksen, I., Quintero, E. A., Harris, C. R., Archibald, A. M., Ribeiro, A. H., Pedregosa, F., and van Mulbregt, P. (2020). SciPy 1.0: fundamental algorithms for scientific computing in Python. Nature Methods, 17(3):261–272. Publisher: Nature Publishing Group.

Vlasits, A., Morrie, R., Tran-Van-Minh, A., Bleckert, A., Gainer, C., DiGregorio, D., and Feller, M. (2016). A Role for Synaptic Input Distribution in a Dendritic Computation of Motion Direction in the Retina. Neuron, 89(6):1317–1330.

Wang, X., Roberts, P. A., Yoshimatsu, T., Lagnado, L., and Baden, T. (2023). Amacrine cells differentially balance zebrafish color circuits in the central and peripheral retina. Cell Reports, 42(2):112055.

Wässle, H. (2004). Parallel processing in the mammalian retina. Nature Reviews Neuroscience, 5(10):747–757.

Yan, W., Laboulaye, M. A., Tran, N. M., Whitney, I. E., Benhar, I., and Sanes, J. R. (2020). Mouse Retinal Cell Atlas: Molecular Identification of over Sixty Amacrine Cell Types. The Journal of Neuroscience, 40(27):5177–5195.

Yatsenko, D., Reimer, J., Ecker, A. S., Walker, E. Y., Sinz, F., Berens, P., Hoenselaar, A., Cotton, R. J., Siapas, A. S., and Tolias, A. S. (2015). DataJoint: managing big scientific data using MATLAB or Python. Pages: 031658 Section: New Results.

Zhao, Z., Klindt, D. A., Maia Chagas, A., Szatko, K. P., Rogerson, L., Protti, D. A., Behrens, C., Dalkara, D., Schubert, T., Bethge, M., Franke, K., Berens, P., Ecker, A. S., and Euler, T. (2020). The temporal structure of the inner retina at a single glance. Scientific Reports, 10(1):4399.

Ölveczky, B. P., Baccus, S. A., and Meister, M. (2003). Segregation of object and background motion in the retina. Nature, 423(6938):401–408.

