## Supplementary Information for "GABAergic amacrine cells balance biased chromatic information in the mouse retina"

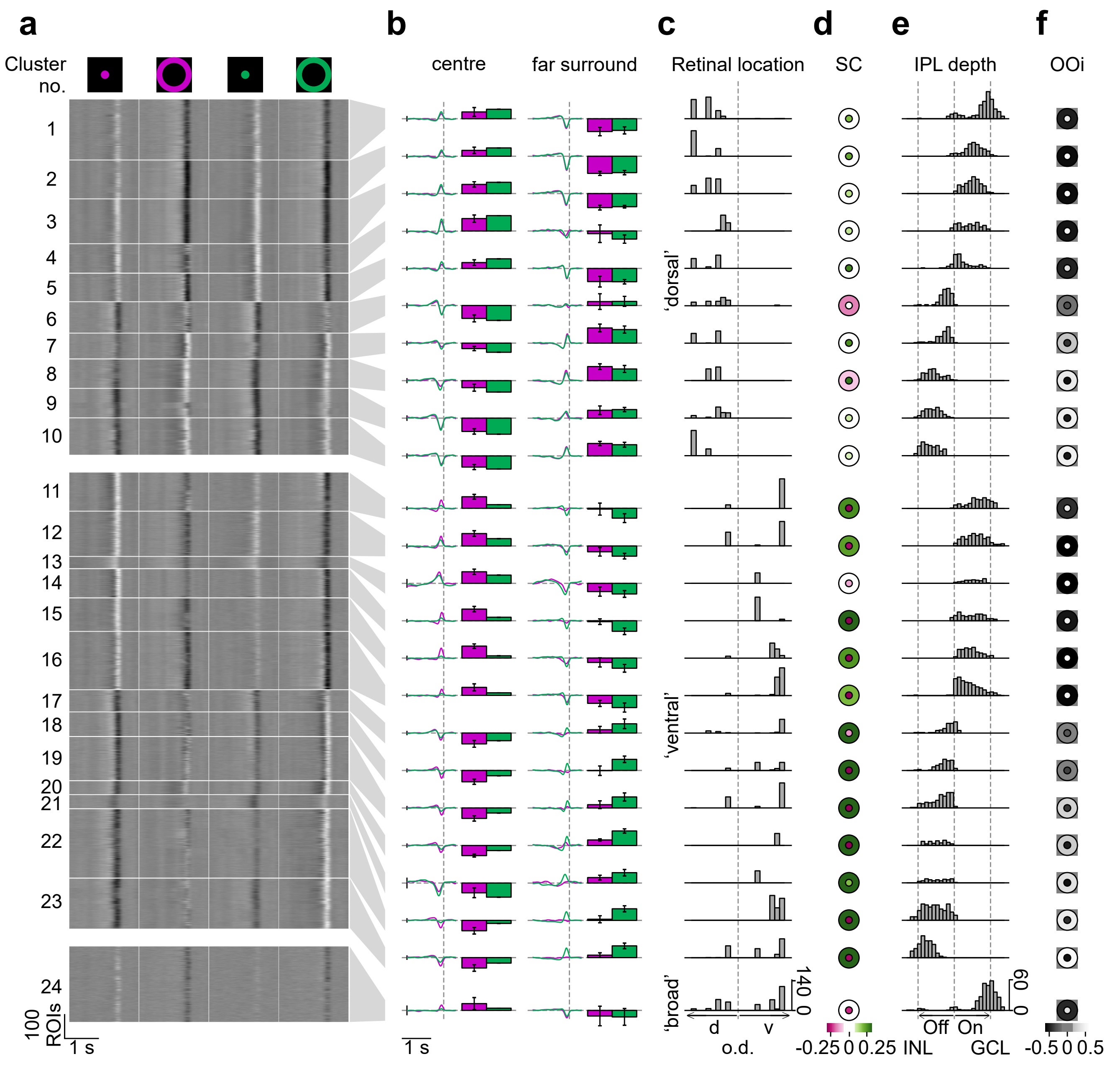


**Fig. Suppl. Fig. 1. Clustering of bipolar cell chromatic responses. (a)** Temporal kernels from 3,589 ROIs' responses to UV and green, centre and far-surround 10 Hz flicker stimulation are extracted and sorted into clusters. Clusters are ordered according to retinal position and average inner plexiform layer (IPL) depth. Scale bars: x, 1 s; y, 100 ROIs. **(b)** Clusters' mean UV/green temporal kernels and colour tuning bars with error bars showing $\pm$s.d., for centre and far-surround. Vertical and horizontal dotted lines indicate response time and baseline, respectively. Scale bars: x, 1 s; y, 0.2 a.u. **(c)** ROI distribution for each cluster along the retina's dorsal (d) – ventral (v) axis. Clusters are sorted into three groups based on retinal location. Clusters made up of 80% or more ROIs from one retinal location are considered "ventral" or "dorsal", other clusters are considered "broad". Dotted line indicates relative optic disc location. **(d)** Representation of clusters' spectral contrast ($SC$) extracted from centre- (inner domain) and far-surround (outer domain) mean temporal kernels. **(e)** ROI distribution for each cluster within the IPL. Left and right dotted lines represent borders with inner nuclear layer (INL) and ganglion cell layer (GCL), respectively. Middle dotted line represents relative border that separates the Off (left) and the On (right) IPL stratum. **(f)** Representation of clusters' On-Off index ($OOi$), similarly as in (d).

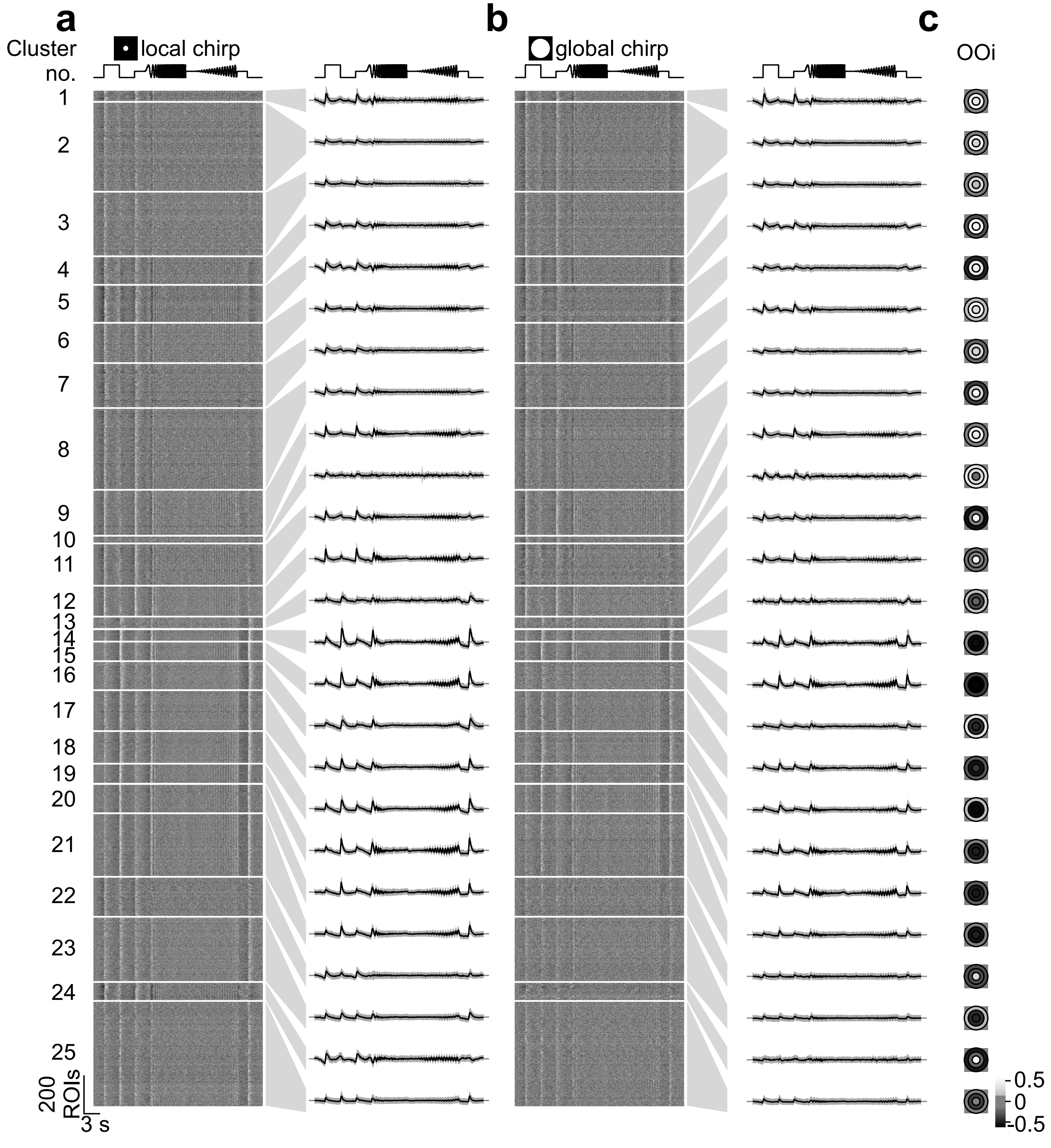


**Fig. Suppl. Fig. 2. Achromatic chirp responses of the 25 AC chromatic clusters. (a)** Left: Responses from 5,378 ROIs to local (100 µm) chirp stimulus, sorted into clusters. Right: Average chirp responses with ±s.d. shading for each cluster. Scale bars: x, 3 s; y, 200 ROIs; horizontal dotted lines: baseline. **(b)** Like (a) but for global (800 µm) chirp stimulus. **(c)** Same as in Fig. 3d: Response polarity as On-Off index ($OOi$) of each cluster's mean chromatic kernels.

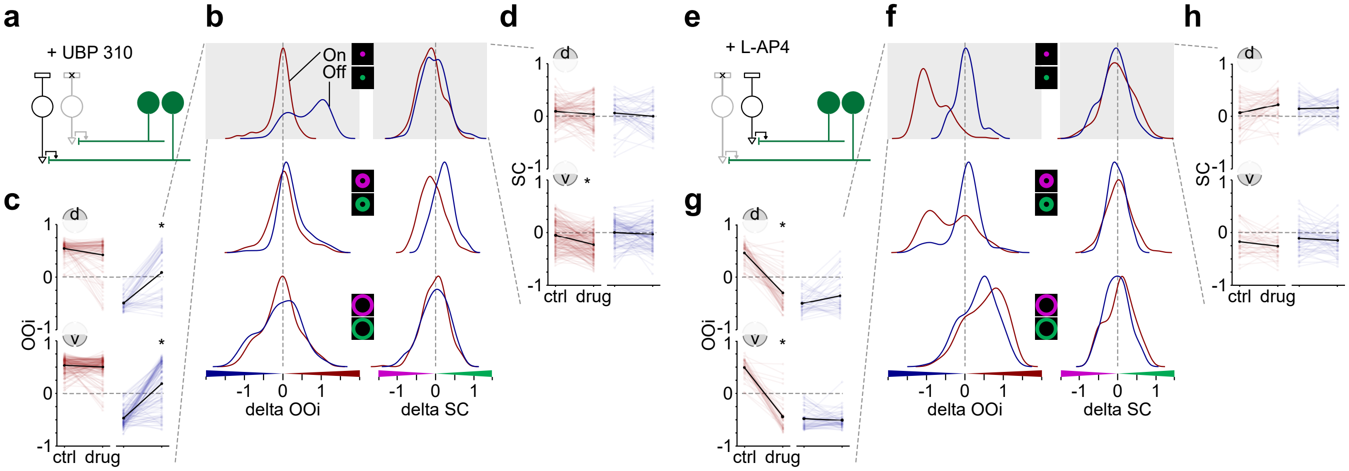


**Fig. Suppl. Fig. 3. Effects of On and Off BC excitation on AC chromatic responses.** **(a)** Schematic showing the effects of blocking the Off excitatory pathway on GABAergic amacrine cell (AC) signalling. UBP 310, (S)-1-(2-Amino-2-carboxyethyl)-3-(2-carboxy-thiophene-3-yl-methyl)-5-methylpyrimidine-2,4-dione (10 µM). Black, shaded black and green cells represent an On bipolar cell (BC), an Off BC and recorded GABAergic ACs, respectively. **(b)** Density plots of the difference in On-Off index (delta $OOi$) and spectral contrast (delta $SC$) between control and drug condition for centre (top), local-surround (middle) and far-surround (bottom) ($n = 445$ ROIs, $n = 8$ scan fields, $n = 4$ mice). We used the $OOi$ of the "centre-control" condition to define On ROIs ($OOi > 0$, $n = 286$ ROIs, red) and Off ROIs ($OOi < 0$, $n = 159$ ROIs, blue). **(c)** Effect of UBP 310 on "center" $OOi$. ROIs are separated in On (red) and Off (blue), as well as "dorsal" (top) and "ventral" (bottom). $*p < 0.05$, "ns", not significant; paired t-test corrected for multiple comparisons. **(d)** Effect of UBP 310 on "center" SC. Statistics same as in (c). **(e)** Schematic showing the effects of blocking the On excitatory pathway on GABAergic AC signalling. L-AP4, L-2-amino-4-phosphonobutyric acid (50 µM). Black, shaded black and green cells represent an Off BC, an On BC, and recorded GABAergic ACs, respectively. **(f-h)** Same as in (b-d) but for evaluating effects of the absence of input to On BCs in the circuit ($n = 200$ (95 On, 105 Off) ROIs, $n = 9$ scan fields, $n = 4$ mice).

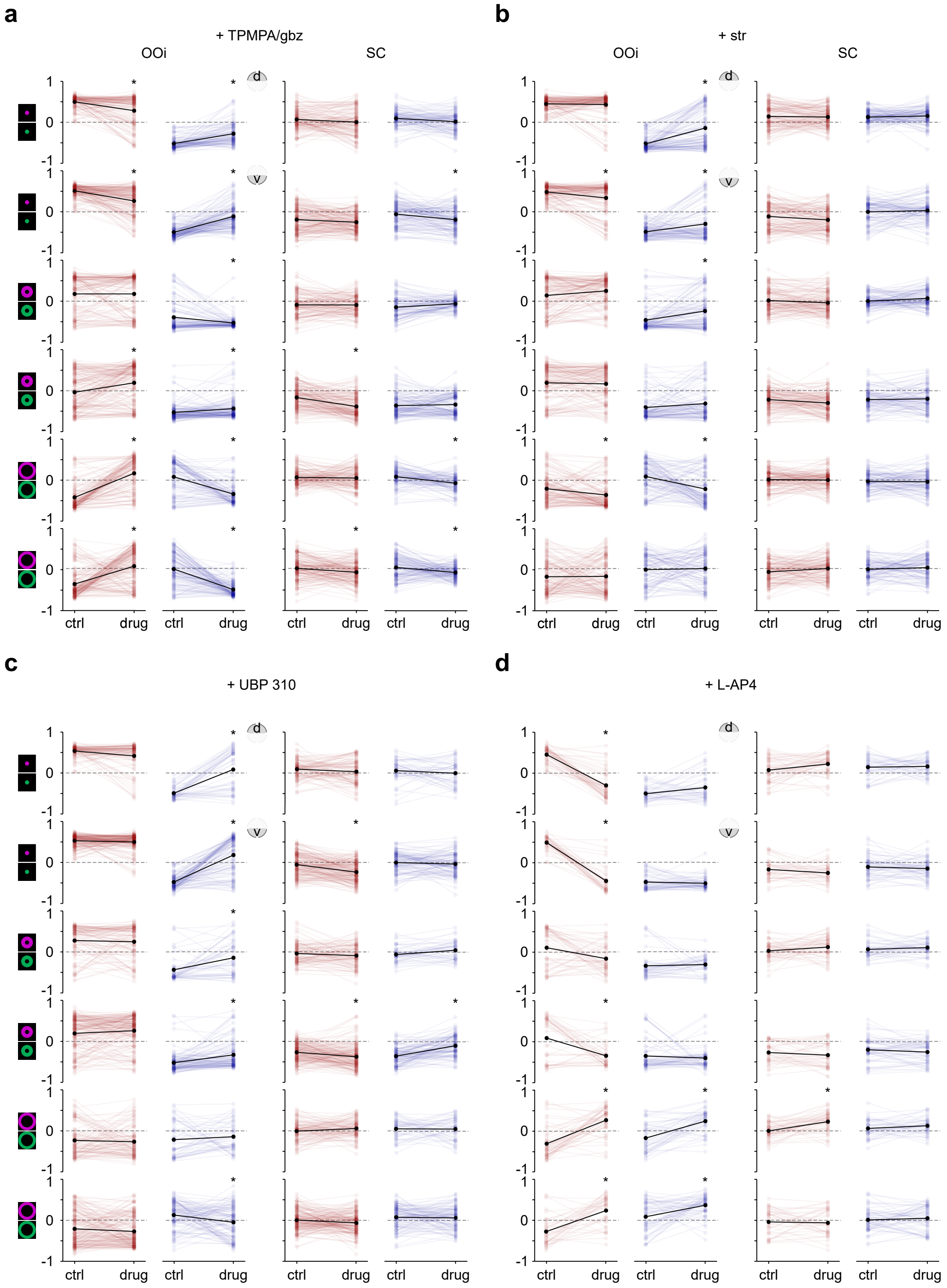


**Fig. Suppl. Fig. 4. Effects of BC excitation and AC inhibition on AC chromatic responses. (a)** Effect of TPMPA/gbz on centre, local-surround, and far-surround On-Off index ($OOi$, left two columns) and spectral contrast ($SC$, right two columns). ROIs are separated in On (red) and Off (blue), as well as "dorsal" and "ventral", as shown in the schematics on the left. $*p < 0.05$, "ns", not significant; paired t-test corrected for multiple comparisons. **(b-d)** Same as in (a) but for strychnine, UBP 310 and L-AP4 application, respectively.

### Model equations

Equations from Schröder et al. (2020) with modifications.

**Photoreceptors.** We modelled individual photoreceptors as a biphasic filter $\kappa^{\text{PR}}(t,\gamma)$ (Baden et al., 2014; Schnapf et al., 1990) with a rise and decay constant ($\tau_r = 0.05$ and $\tau_d = 0.05$, respectively) and phase parameters $\phi$ and $\tau_{\text{phase}}$ ($\phi = -\pi/7$ and $\tau_{\text{phase}} = 100$). The photoreceptor filter had one free parameter $\gamma$ to account for temporal kinetics (Schröder et al., 2019) and was normalised to have norm one. The light input $l$ was processed by photoreceptors resulting in the photoreceptor output $\text{PR}^{\text{out}}$. The total input to a bipolar cell $\text{BC}^{\text{in}}$ was given by a weighted sum of the output of the six photoreceptors (UV and green; centre, local and far surround). The PRs summarise all temporal processing upstream of the BC terminal.

$$\kappa^{\text{PR}}(t,\gamma) = \frac{-\left(\frac{t}{\gamma\tau_r}\right)^3}{1+\frac{t}{\gamma\tau_r}} \cdot \exp\left(-\left(\frac{t}{\gamma\tau_d}\right)^2\right) \cdot \cos\left(\frac{2\pi t}{\gamma\phi} + \tau_{\text{phase}}\right)$$

$$\text{PR}^{\text{out}}(t) = \int_{\tau=0}^{T} l(t-\tau) \cdot \kappa^{\text{PR}}(\tau)d\tau.$$

$$\text{BC}^{\text{in}}(t) = \sum_{i=1}^{6} w_i^{\text{PR BC}} \cdot \text{PR}_{\text{i}}^{\text{out}}$$

**Bipolar cells.** The input to bipolar cells was modulated by the inhibitory feedback of amacrine cells (see below) $\text{fb}_{\text{AC}}$ and then subjected to a non-linearity (see below) to obtain the vesicle release probability $p$. The vesicles released were based on the current release probability $p$ and the vesicles available in the readily-releasable pool (RRP). Vesicles were moved from the cytoplasm to the intermediate pool (IP) at rate $k2$ and from the IP to the RRP at rate $k1$ to replenish the RRP.

$$p(t) = \sigma\left(\text{BC}^{\text{in}}(t) - \text{fb}_{\text{AC}}(t)\right).$$

$$\text{release} = p(t) \cdot \text{RRP}, \qquad \text{RRP}_{\text{refill}} = k_1 \cdot \text{IP}, \qquad \text{IP}_{\text{refill}} = k_2,$$

**Non-linearity.** The sigmoid we used as the non-linearity had an offset $x_0$ and slope $k$ as free parameters.

$$\sigma(x) = \frac{1}{1+\exp(-k(x-x_0))}.$$

**Amacrine cells.** Each amacrine cell was modelled using a double exponential kernel $\kappa^{\text{AC}}$ with a rise and decay constant, $\tau_r$ and $\tau_d$, respectively. Amacrine cells received a weighted input from all 13 bipolar cells and to obtain their output, we applied a non-linearity (see above). The feedback signal to bipolar cells $\text{fb}_{\text{AC}}$ was obtained by first applying amacrine cell to amacrine cell inhibition to the output signal $\text{AC}^{\text{out}}$. A weighted sum of the 25 amacrine cell signals then constituted the feedback to bipolar cells.

$$\kappa^{\text{AC}}(t) = \exp\left(\frac{-t}{\tau_d}\right) - \exp\left(\frac{-t\cdot(\tau_d+\tau_r)}{\tau_d \cdot \tau_r}\right).$$

$$\text{AC}^{\text{out}}(t) = \sigma\left(\int_{\tau=0}^{T}\left(\sum_{j=1}^{13} w_j^{\text{BC AC}} \cdot \text{BC}_j^{\text{out}}(t-\tau)\right) \cdot \kappa^{\text{AC}}(\tau)d\tau\right).$$

$$\text{fb}_{\text{AC}}(t) = \sum_{j=1}^{25} w_j^{\text{AC BC}} \cdot \left(\text{AC}_j^{\text{out}}(t) - \sum_{n=1}^{25} w_{nj}^{\text{AC AC}} \cdot \text{AC}_n^{\text{out}}(t)\right).$$
